# Structural insights into distinct mechanisms of RNA polymerase II and III recruitment to snRNA promoters

**DOI:** 10.1101/2024.09.10.612236

**Authors:** Syed Zawar Shah, Thomas N. Perry, Andrea Graziadei, Valentina Cecatiello, Thangavelu Kaliyappan, Agata D. Misiaszek, Christoph W. Müller, Ewan P. Ramsay, Alessandro Vannini

## Abstract

RNA polymerase III (Pol III) is specialized in the transcription of short, essential RNAs, including the U6 small nuclear RNAs (snRNAs). At U6 snRNA genes, Pol III is recruited by the snRNA Activating Protein Complex (SNAPc) forming, together with a Brf2-containing TFIIIB complex, a transcriptionally competent pre-initiation complex (PIC). Additionally, SNAPc is responsible for the recruitment of Pol II at the remaining snRNAs genes (U1, 2, 4 and 5), representing a unique example of a multi subunit transcription factor shared among different RNA Polymerases. The mechanism of SNAPc cross-polymerase engagement and the role of the SNAPC2 and SNAPC5 subunits in transcription remain poorly defined. Here, we present cryo-EM structures of the full-length SNAPc-containing Pol III PIC assembled on the U6 snRNA promoter in the open and melting states at 3.2-4.2Å resolution. Comparative structural analysis revealed unexpected differences with the yeast PIC and revealed the molecular basis of selective and structurally distinct SNAPc engagement within Pol III and Pol II PICs. Harnessing crosslinking mass spectrometry, we also localize the SNAPC2 and SNAPC5 subunits in proximity to the bound promoter DNA, expanding upon existing descriptions of snRNA Pol III PIC structure.

## Introduction

Transcription of the eukaryotic genome relies on three multi-subunit RNA polymerase (Pol) complexes: Pol I, II and III. Pol III specialises in the transcription of small, non-translated RNA species encompassing the 5S rRNA, the entire pool of cellular tRNAs and many essential RNA products such as the U6 snRNA, 7SL, 7SK, RPPH1 and RMRP^1^. During transcription, Pol III engages with a host of general transcription factors (GTFs), to form a pre-initiation complex (PIC) which facilitates transcription initiation and represents a major regulatory step in the transcription cycle^2^.

Pol III transcribes a diverse set of RNAs and, accordingly, relies on a diverse set of promoters and associated GTFs which form multiple different PIC complexes at Pol III genes^3^. Canonical class I and II genes are characterized by promoter sequences encompassed within the coding region, recognized directly or indirectly by the TFIIIC complex, in turn allowing recruitment of RNA Pol III via a Brf1-containing TFIIIB complex.

Found specifically in vertebrates, the class III promoter, which directs the transcription of a small subset of approximately 20 genes, including the U6 snRNA, 7SL and RPPH1^4,5^, represents an exception to the general architecture of Pol III promoters. The class III promoter is characterized by a unique architecture, with 3 conserved sequences found upstream of the transcriptional start site (TSS) comprising a TATA box (around 20-25 base pairs upstream), the proximal sequence element (PSE, found between 40-70 base pairs from the TSS) and the distal sequence element (DSE, located approximately 200 bp upstream)^6,7^. The PSE directs the first stage of class III transcription through recruitment of the hetero-pentameric snRNA activating protein complex (SNAPc), also known as the PTF (PSE transcription factor) or PBP (PSE binding protein)^8–12^. SNAPc consists of 5 subunits known as SNAPC1 (SNAP43), SNAPC2 (SNAP45), SNAPC3 (SNAP50), SNAPC4 (SNAP190) and SNAPC5 (SNAP19)^12–17^ and is organized around a central core consisting of the SNAPC4 N-terminal region, SNAPC3 and SNAPC1 which represent the minimal assembly (termed miniSNAPc) required for RNA Pol III transcription^18^, with SNAPC4 and SNAPC3 constituting the major DNA binding site^13,14,19^. In contrast, while apparently dispensable for transcription, SNAPC5 and SNAPC2 are required to stabilize the assembly and overcome SNAPC4 auto-inhibition of DNA binding^14,18,20^. Recruitment of SNAPc is facilitated via interactions with Oct-1 and ZNF143 bound at the DSE^9,21,22^, which are juxtaposed via a positioned nucleosome found between the PSE and DSE^23,24^. Following stable recruitment, SNAPc can, in turn, facilitate recruitment of an alternative TFIIIB heterotrimeric transcription factor consisting of the TATA box binding protein (TBP), B-double prime (Bdp1) and a variant of Brf1, termed Brf2, to the downstream TATA box, allowing subsequently for Pol III recruitment and transcription initiation^25–29^. Despite the limited pool of genes transcribed by the class III promoter, regulation of this process is important, with altered class III transcription due to overexpression of Brf2 associated with cancer^30^ and SNAPc mutations which reduce snRNA transcription identified in patients with neurodevelopmental disorders^31^. Furthermore, the RNA Pol III U6 snRNA promoter is widely used to drive transcription of single guide RNA in most CRISPR/Cas9 systems^32^.

Additionally, understanding the molecular basis of SNAPc as a general transcription factor to direct Pol III transcription at class III promoters is particularly relevant taking into consideration the involvement of SNAPc in Pol II-mediated transcription of the remaining snRNA genes^33,34^. SNAPc represents a unique case of a multi-subunit general transcription factor capable of engaging with two distinct transcription machineries while preserving promoter specificity. Indeed, the polymerase recruited downstream of a bound SNAPc complex is wholly determined by the presence or absence of a TATA box, which designates the loci for either Pol III (such as for U6 snRNA) or Pol II (such as for U1, U2, U4 and U5 snRNA) transcription, respectively^35^. Despite structures of miniSNAPc bound to the U6 promoter^36^ and miniSNAPc-containing Pol III^37^ and Pol II^38^ PICs being reported, a mechanistic understanding of how SNAPc is able to faithfully recruit the correct polymerase at their cognate snRNA genes is currently not understood. Here, we present cryo-EM structures of the U6 snRNA Pol III open complex PIC assembled with full length and mini SNAPc at a resolution range of 3.2 – 4.2Å. Our results provide detailed insight into the assembly of the SNAPc-TFIIIB complex with Pol III at type III promoters and allow for direct comparison of Pol III PIC structures to those previously solved for *Saccharomyces cerevisiae*^39–41^ revealing unexpected differences in the conformational changes observed in the polymerase upon PIC formation and transcription initiation. Employing an integrative approach, the SNAPC2 and SNAPC5 subunits were localized, providing the first structural analysis of these subunits in the SNAPc complex. Comparative structural analysis also revealed the structural basis of the specific recruitment of Brf2 to class III promoters and, finally, identified a novel double-sided interaction motif in SNAPC4 which confers the unique ability of SNAPc to engage in both Pol II and Pol III PICs while retaining promoter specificity.

## Results

### Cryo-EM Visualization of the human class III PIC

To visualize the structure of the class III PIC, recombinant full-length SNAPc (SNAPc^FL^), Brf2, TBP, Bdp1 were purified as previously described^5,29,38^ and assembled with purified endogenous human Pol III on the U6-2 class III scaffold (Extended Data Figure 1). The reconstituted PIC (SNAPc^FL^-PIC) complex was subsequently purified and simultaneously cross-linked via GraFix to produce PIC complexes which were resolved by cryo-EM (Figure 1; Extended Data Figures 1-3). Three-dimensional classification of the dataset observed three distinct states for the PIC, two of which corresponded to an open complex (OC), where the DNA was spontaneously opened in the polymerase active site. Of those, one displayed a more ‘closed’ clamp engaged with the DNA with a channel width of 19.9 Å (OC1) and the other possessing a ‘open’ clamp with a channel width of 21.6 Å (OC2). The third observed state represented an intermediate melting complex (MC) with a significantly wider RPC1 clamp (with channel width of 32.2 Å) and a disordered downstream DNA, as observed previously^37^ (Extended Data Figures 2-4). In each state, the density corresponding to the upstream bound TFIIIB:SNAPc^FL^ module was poor, and so was subjected to a masked classification focused on this region. The resulting classes produced multiple states for each OC and MC in which SNAPc^FL^ made multiple contacts (fully engaged) or only a single contact (partially engaged) with the downstream TFIIIB and Pol III densities (Extended Data Figures 2-4). Subsequent refinement resulted in three reconstructions for the SNAPc^FL^-PIC corresponding to fully engaged TFIIIB:SNAPc with a Pol III closed clamp (OC1^FL^, at an overall resolution of 3.26Å), a partially-engaged TFIIIB:SNAPc with a Pol III open clamp (OC2^FL^, at an overall resolution of 3.4Å) and MC^FL^ with a fully engaged TFIIIB:SNAPc module (resolved at an overall resolution of 3.51Å) (Table 1). Comparison between OC1^FL^ and OC2^FL^ densities showed no apparent co-ordination between the observed clamp and TFIIIB:SNAPc movement, suggesting that these are independent motions present in the PIC complex.

**Figure 1.**
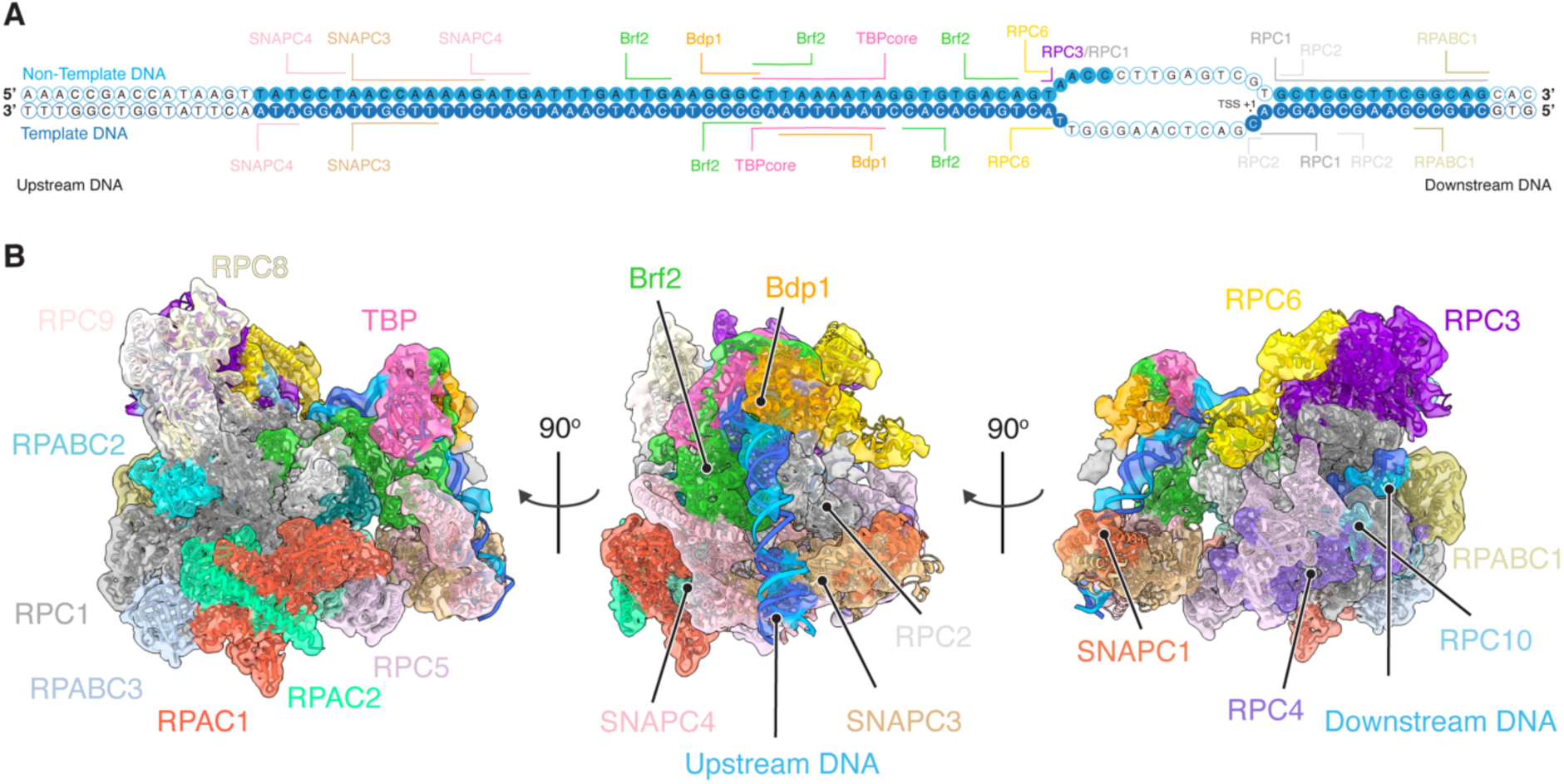
Cryo-EM structure of the human Pol III class III PIC. (a) DNA nucleotides modelled in the OC^FL^ structure are denoted as solid circles, with template and non-template strands indicated. All nucleotides are assigned according to the TSS in the human U6-2 gene. Protein-DNA interactions are labelled with the interacting proteins identified. (b) OC^FL^ locally filtered cryo-EM density map with fitted structural model shown in ribbon representation. The Cryo-EM density is coloured according to the fitted subunit.

**Table 1.** Cryo-EM data collection, refinement and validation statistics.

| Structure Name | OC1<sup>FL</sup> | OC2<sup>FL</sup> | MC<sup>FL</sup> | OC1<sup>mini</sup> | OC2<sup>mini</sup> |
| --- | --- | --- | --- | --- | --- |
| **EMDB Identifier** | EMDB-50730 | EMDB-50731 | EMDB-50732 | EMDB-50733 | EMDB-50734 |
| **PDB Identifier** | PDB-9FSO | PDB-9FSP | PDB-9FSQ | PDB-9FSR | PDB-9FSS |
| **Data Collection and Processing** |  |  |  |  |  |
| **Magnification** | 130 000x | 130 000x | 130 000x | 130 000x | 130 000x |
| **Voltage (KeV)** | 300 | 300 | 300 | 300 | 300 |
| **Total Electron Exposure (e-/Å<sup>2</sup>)** | 50 | 50 | 50 | 50 | 50 |
| **Dose Rate (e-/pix/s)** | 8.59 | 8.59 | 8.59 | 7.29 | 7.29 |
| **Pixel Size (Å/pix)** | 0.955 | 0.955 | 0.955 | 0.955 | 0.955 |
| **Defocus Range (µm)** | -0.75 to -1.75 | -0.75 to -1.75 | -0.75 to -1.75 | -1 to -2.5 | -1 to -2.5 |
| **Symmetry Operator** | C1 | C1 | C1 | C1 | C1 |
| **No. Micrographs Collected** | 17291 | 17291 | 17291 | 18667 | 18667 |
| **Camera Utilised** | Falcon4i | Falcon4i | Falcon4i | Falcon4i | Falcon4i |
| **Energy Filter Utilised** | Selectris X | Selectris X | Selectris X | Selectris X | Selectris X |
| **Energy Slit Size (eV)** | 10 | 10 | 10 | 10 | 10 |
| **Reconstruction** |  |  |  |  |  |
| **Software Utilised** | CryoSPARC (v4.3.1), RELION-4 | CryoSPARC (v4.3.1), RELION-4 | CryoSPARC (v4.3.1), RELION-4 | CryoSPARC (v4.3.1), RELION-4 | CryoSPARC (v4.3.1), RELION-4 |
| **Initial Particle Images (no.)** | 1252298 | 1252298 | 1252298 | 559728 | 559728 |
| **Final Particle Images (no.)** | 15293 | 10808 | 9269 | 15386 | 13256 |
| **Map Resolution (Å)** | 3.26 | 3.4 | 3.51 | 3.76 | 4.14 |
| **FSC Threshold** | 0.143 | 0.143 | 0.143 | 0.143 | 0.143 |
| **Map Sharpening *B* Factor (Å<sup>2</sup>)** | -53.7 | -46.8 | -39.9 | -40.8 | -53.1 |
| **Model Composition** |  |  |  |  |  |
| **Non-hydrogen atoms (no.)** | 56897 | 56808 | 56541 | 56344 | 56344 |
| **Protein Residues** | 6762 | 6762 | 6762 | 6717 | 6717 |
| **Nucleotide Residues** | 139 | 136 | 123 | 132 | 132 |
| **Ligands (Symbol:No.)** | SF4:1, Zn:9, Mg:1 | SF4:1, Zn:9, Mg:1 | SF4:1, Zn:9, Mg:1 | SF4:1, Zn:9, Mg:1 | SF4:1, Zn:9, Mg:1 |
| **Refinement** |  |  |  |  |  |
| **Map CC** | 0.73 | 0.74 | 0.75 | 0.65 | 0.6 |
| **Map vs. Model FSC (Å)** | 3.5 | 3.5 | 3.7 | 4.0 | 6.8 |
| **Map vs. Model FSC Threshold** | 0.5 | 0.5 | 0.5 | 0.5 | 0.5 |
| **r.m.s. deviations** |  |  |  |  |  |
| **Bond Lengths (Å)** | 0.005 | 0.007 | 0.009 | 0.01 | 0.005 |
| **Bond Angles** | 1.198 | 1.221 | 1.292 | 1.284 | 1.083 |
| **B factors (Å<sup>2</sup>)** |  |  |  |  |  |
| **Protein** | 51.97 | 86.77 | 94.81 | 17.06 | 22.72 |
| **Nucleic Acid** | 117.91 | 183.85 | 191.98 | 42.43 | 51.59 |
| **Ligand** | 82.43 | 108.42 | 118.69 | 19.89 | 25.94 |
| **Validation** |  |  |  |  |  |
| **MolProbity Score** | 1.84 | 1.79 | 1.95 | 1.94 | 1.92 |
| **Clashscore (all atom)** | 9.44 | 8.11 | 10.88 | 11.21 | 10.82 |
| **Poor Rotamers (%)** | 0.64 | 0.7 | 0.62 | 0.52 | 0.64 |
| **Ramachandran Plot** |  |  |  |  |  |
| **Favoured (%)** | 95.03 | 94.91 | 94.22 | 94.53 | 94.71 |
| **Allowed (%)** | 4.7 | 4.76 | 5.54 | 5.37 | 5.19 |
| **Disallowed (%)** | 0.27 | 0.33 | 0.24 | 0.11 | 0.11 |

The overall structure revealed a SNAPc complex bound to the upstream PSE engaged with the Brf2 subunit of TFIIIB complex to facilitate TFIIIB:SNAPc assembly (Figure 1). TBP binding to the TATA box induced a sharp bend in the DNA which feeds the promoter DNA into the polymerase between the RPC2 protrusion and RPC1 clamp helices (Extended Data Figure 4G). In the PIC, the nearby flexible RPC6 winged-helix domains 1 and 2 (WH1 and WH2) become stabilized, forming a ‘channel’ which stabilizes the DNA prior to entry into the active site (Extended Data Figure 4H). Consistent with the role of the Brf2 residue C361 in redox regulation^5^ of PIC assembly, it is found solvent exposed on the periphery of the complex with a calculated solvent accessible surface area of 330Å^2^, ensuring this regulation in the PIC context (Extended Data Figure 4I). In each reconstruction, spontaneous DNA opening in the active site produces a highly disordered DNA bubble between -12 and +3 which is partially visible in the map (Extended Data Figure 4J). Comparison of models for the OC1, OC2 and MC states also observed a highly similar structure for the TFIIIB:SNAPc region, suggesting that this represents a rigid motion of the upstream transcription factors with respect to the polymerase (Extended Data Table 1). Despite PIC assembly with the full-length protein, the density for the SNAPc^FL^ complex corresponded to a core assembly of the SNAPC4 N-terminus lacking the MyB DNA-binding domain (residues 142-383), SNAPC3 and the N-terminal half of SNAPC1 (residues 1-141), with the SNAPC2 and SNAPC5 subunits not observed (Extended Data Figure 4K), consistent with their dispensable role in DNA binding^17,42^. This closely resembled the previously described miniSNAPc, designated the minimal SNAPc machinery required to facilitate PIC formation and Pol III transcription^18^.

For a direct comparison, we additionally imaged the class III PIC assembled with SNAPc^mini^ consisting of SNAPC4 (1-505), SNAPC3 and SNAPC1 (1-268) using Cryo-EM. Three-dimensional classification was carried out as previously, yielding exclusively ‘closed-clamp’ and ‘open-clamp’ OCs (Extended Data Figure 5). Only reconstructions for the ‘closed’ OC produced TFIIIB:SNAPc^mini^ density of sufficient quality for TFIIIB:SNAPc^mini^ masked classification and refinement, producing two OC structures (OC1^mini^ resolved at 3.76Å resolution and OC2^mini^ resolved at 4.1Å resolution) which corresponded to SNAPc^mini^ fully and partially engaged TFIIIB:SNAPc OC structures, respectively (Extended Data Figures 5-6). In each instance, both SNAPc^FL^ and SNAPc^mini^ structures were very similar (Extended Data Table 1) and each displayed a similar fold to the miniSNAPc-DNA complex resolved in isolation (Extended Data Table 1; Extended Data Figure 6I-K). Together, these results suggest that the SNAPc core is unaffected by the presence of the mobile SNAPC2 and SNAPC5 subunits and is well defined, presenting PIC assembly-competent interfaces without any requirement for complex remodelling. As all OC structures are highly similar and represent equivalent functional states, the OC1^FL^ structure, which displayed the highest quality reconstruction, was selected and will be hereafter referred to as OC^FL^. All the following presented structural analysis was carried out with this model unless otherwise specified.

### Brf2 uses a conserved interaction to facilitate transcription initiation

Structural comparison of the human (*H.s.*) class III PIC to the *Saccharomyces cerevisiae (S.c.)* PIC formed on the SNR6 promoter revealed that Brf2 shared a common mechanism of Pol III interaction with the *S.c.* Brf1. Sequence and structural alignment of the N-terminal Zn ribbon/B-reader/B-linker region for both *S.c.* Brf1^41^ and *H.s.* Brf2 displayed a high level of conservation and structural similarity (Figure 2A-B). Alignment of both the human and *S.c.* PIC structures identified an equivalent docking position for Brf1 and Brf2 on Pol III between the human and yeast structures, with the Pol III region involved presenting highly conserved interfaces (Figure 2C; Extended Data Figure 7A), demonstrating an equivalent mode of interaction irrespective of species and promoter type.

**Figure 2.**
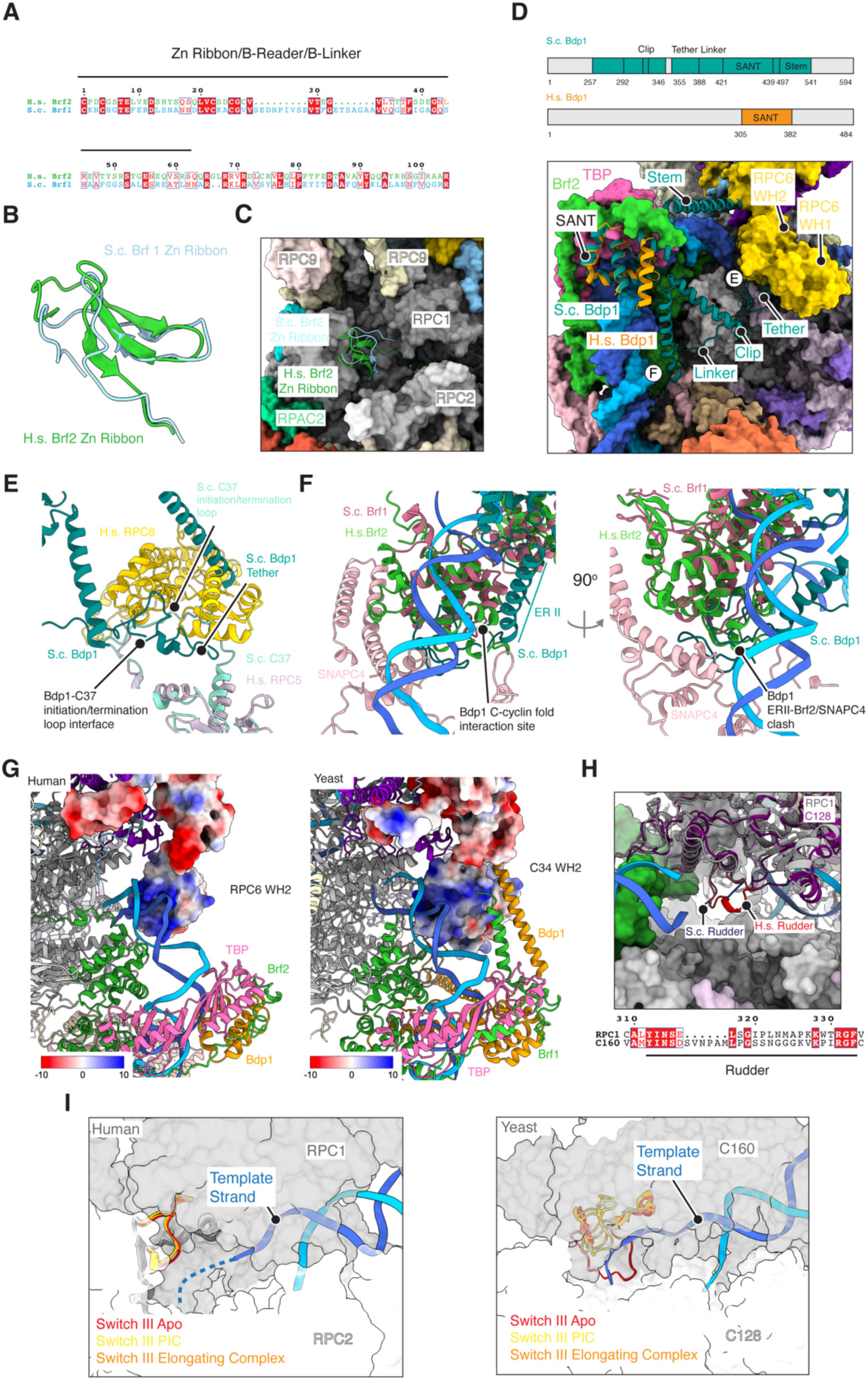
Comparison of human and yeast snRNA PIC complexes. (a) Sequence alignment of the N-terminal region of human Brf2 and yeast Brf1, comprising the Zn ribbon/B-linker/B-reader region. Conserved residues are highlighted in red, while those with sequence similarities are outlined in blue. (b) Structural alignment of the human (H.s.) Brf2 (green) and yeast (S.c.) Brf1 (PDB: 6eu0, blue) Zn ribbon structures from their respective PICs. (c) Alignment of the human (H.s.) Brf2 and yeast (S.c.) Brf1 (PDB: 6eu0) Zn ribbon structures showing their relative positions with respect to Pol III. Shown in surface representation is the human Pol III structure. (d) A schematic representation of the yeast (S.c.) and human (H.s.) Bdp1 present in their respective PICs. Built regions are shown in colour (*above*) with structural alignment between human (orange) and yeast (teal) Bdp1 shown in the context of the human PIC (*below*). Altered Bdp1 interfaces are highlighted with (e) and (f). (e) Structural alignment of human RPC5 and yeast C37, with the yeast Bdp1 (teal) superimposed highlighting the loss of initiation/termination loop structure in RPC5 and interaction interface with Bdp1 in the human PIC. (f) Structural alignment of human (H.s.) Brf2 (green) and yeast (S.c.) Brf1 (pink), with the yeast Bdp1 (teal) superimposed, displaying an altered interaction interface between Bdp1 and Brf2 and clashes with both Brf2 and SNAPC4 in the human PIC. (g) Comparison of the upstream promoter DNA entry into the human (U6_2, *left*) and yeast (PDB:6eu0, SNR6, *right*) polymerase. In both, human RPC6 (*left*) and yeast C34 (*right*) WH1 and 2 domains are shown in a surface representation rendered according to colombic surface potential. Shown in both cases are the basic surface patches presented to the incoming DNA. (h) Structural alignment of the human (red) and yeast (PDB: 6eu0, blue) rudder shown in the context of the human PIC, together with the sequence alignment (*below*). (i) Structural alignment of the human apo Pol III, elongating complex and PIC (PDB: 7a6h and 7ae1) (*left*) and yeast apo Pol III, elongating complex and PIC (PDB: 6eu3, 6eu1 and 6eu0) (*right*). In each case the position of the switch III loop is compared relative to the template strand of the open DNA bubble.

In contrast, the Brf1 and Brf2-associated factor Bdp1 displays significant structural differences between the human and yeast PICs. Both *H.s.* and *S.c*. Bdp1 form a conserved SANT domain which facilitates interactions with TFIIIB (Figure 2D). However, in the yeast PIC, Bdp1 undergoes a significant conformational change from a disordered to a largely ordered structure. The N-terminal ‘tether’ domain folds to engage the C37 ‘initiation/termination loop’ and stabilizes the WH1 and WH2 domains of the *S.c.* RPC6 homologue C34^39,41^. C-terminal to this, the ‘clip’ domain (also previously identified as the ERII region, encompassing residues 269-312 in yeast)^39,43^ folds to form contacts with the C-cyclin domain of Brf1^41^. No such transition is observed for the human system, with both regions N-and C-terminal to the SANT domain remaining disordered upon PIC formation (Figure 2D). Alignment of human and yeast structures revealed a loss of the ordered initiation-termination loop in human RPC5 (Figure 2E) together with an altered structure of the ‘clip’ domain binding interface in Brf2, with significant clashes also observed between the N-terminus of Bdp1 and Brf2 and SNAPC4 in the human U6 PIC structure (Figure 2F). Removal of these stabilising interactions likely leads to the observed loss of Bdp1 structure in the human system. Given the established role of these regions in transcription initiation^43^, this suggests a potential difference in the mechanism of transcription initiation at human and yeast U6 genes. Despite the loss of Bdp1 structure, the human RPC6 WH1 and 2 adopt a highly similar conformation to the yeast, which was unexpected in the absence of an ordered Bdp1 stem and tether. Nevertheless, a comparison between both structures revealed a basic surface patch presented by RPC6 WH2 in both human and yeast structures (Figure 2G), suggesting that the interaction with the upstream DNA is sufficient to stablise RPC6 WH1 and 2 upon formation of the PIC.

The structural comparison also discerned differences within the polymerase active site. Whilst most functional regions including fork loops I and II, switch II, the bridge helix and the trigger loop were identical, as suggested by their sequence conservation (Extended Data Figure 7B-C), differences were observed in the rudder and switch III loop. Comparison of *H.s.* and *S.c.* rudder sequences revealed a divergence between both regions which was also observed in the structure, with the human rudder adopting a highly similar folded structure in apo, elongating and PIC engaged states in contrast to the yeast polymerase, where this region is disordered in the PIC (Figure 2H) and folds upon transition to the elongating complex (EC)^41^. The rudder forms stabilizing interactions with the RNA-DNA hybrid in the active site, suggesting a difference in transcription initiation^44^. Differences were also observed in the switch III loop despite high sequence conservation. Importantly, *S.c.* Brf1 N-terminal domain is critical for full expansion of the initial transcription bubble around position -9 via an allosteric mechanism through direct contact with the Switch III loop^41,45^. Indeed, *S.c.* Brf1 Zn-ribbon reconfigures the switch III loop, preventing the occlusion of the template strand from the active site and allowing for the stabilisation of the transcription bubble^41^. However, the human switch III loop is constitutively open in all states^46^ (Figure 2I) suggesting that human and yeast Pol III-mediated DNA opening, and therefore transcriptional initiation, may be differentially regulated.

### SNAPc structure within the Class III PIC

Resolution of the SNAPc ‘core’ structure in the OC context observed a similar structure to the PSE-bound miniSNAPc complex^36^ (Extended Data Table 1). In the SNAPc core assembly, SNAPC3 represents a central structural unit around which the entire SNAPc complex is assembled (Figure 3A). The first 5 ⍺-helices in the N-terminus of SNAPC3 (the ‘lasso’ domain)^36^ together with an anti-parallel β-sheet of the ubiquitin like domain (ULD) completely enclose the N-terminus of SNAPC1 (Figure 3B). SNAPC4 assembles on the opposing face of SNAPC3, forming multiple contacts which enclose the ‘wedge’ region (Figure 3C-D). The N-terminal region (M142-G158) of SNAPC4 inserts into a groove formed by the SNAPC3 ULD (Figure 3B). Additional hydrophobic contacts are formed by the R_h_ and R_a_ repeats of the SNAPC4 MyB domain on the opposing surface (Figure 3C). In the Pol II PIC, SNAPc was observed to contain two protrusions, known as wing 1 (formed by SNAPC4) and wing 2 (a helical bundle formed by SNAPC1, SNAPC4 and SNAPC5). In our structure, only wing 1 was observed with wing 2 not present (Figure 3A). This is likely due to the loss of a stabilizing interaction between TFIIB and wing 2 observed in the Pol II PIC, rendering this region flexible in the Pol III PIC and so absent from the OC^FL^ map. Indeed, the miniSNAPc-PSE structure determined previously observed a similar loss of both the wing 1 and wing 2 structures^36^, suggesting that these structures are only stabilized upon binding to cognate transcription factors.

**Figure 3.**
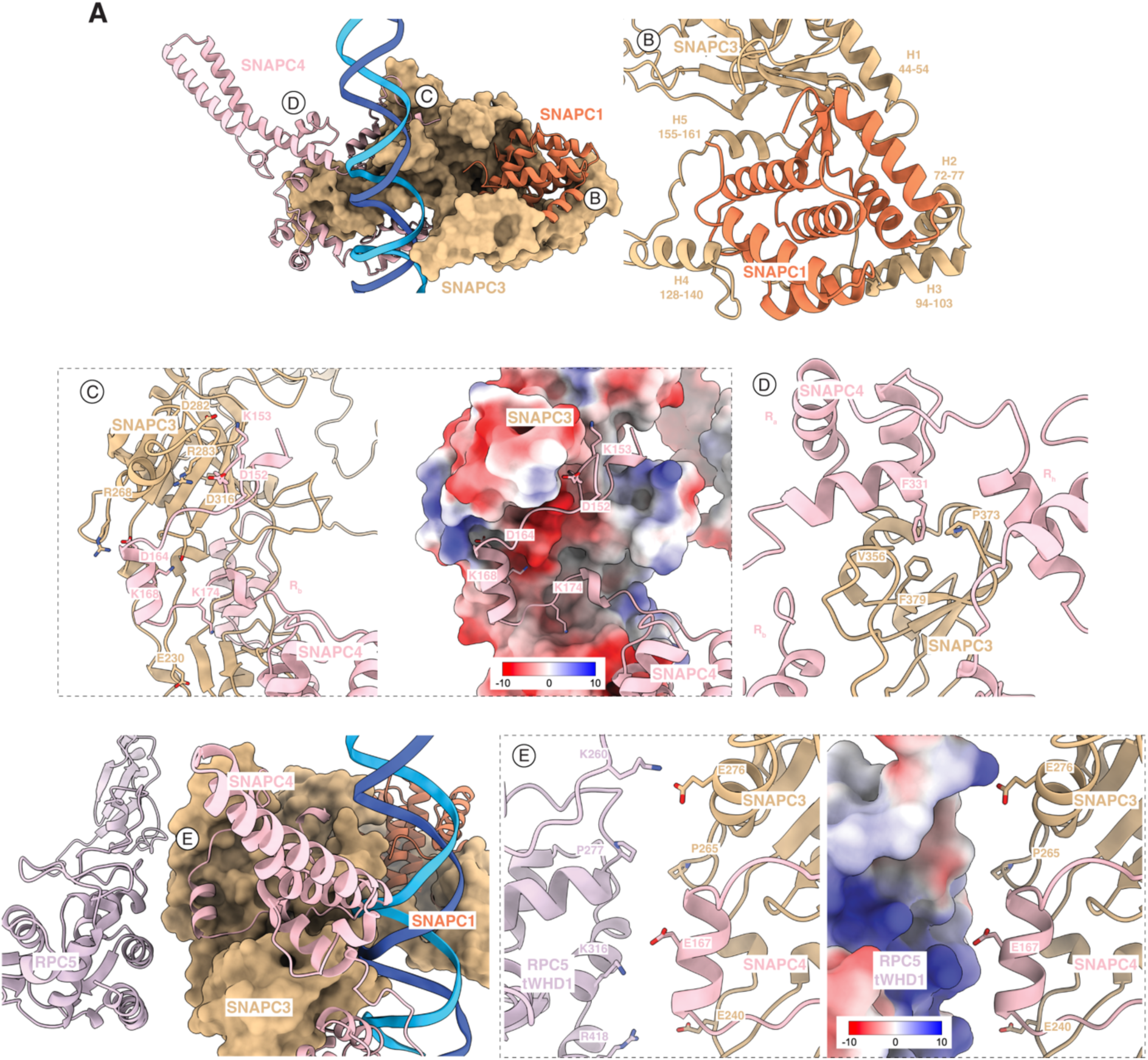
Architecture of SNAPc in the Pol III PIC. (a) The overall structure of SNAPc in the Pol III PIC. SNAPC3 is shown in surface representation with both SNAPC1 and SNAPC4 shown as ribbons. Interfaces between SNAPC3 and other SNAPc subunits are highlighted (d-d) with interacting residues labelled. (e) The interface between SNAPC3 and RPC5 tWHD1, with interacting residues highlighted.

In the OC^FL^, SNAPC3 and SNAPC4 were also observed to contact the RPC5 C-terminal tandem winged helix domain 1 (tWHD1) in Pol III. The acidic residues E167 of SNAPC4 and E240, E266 and E276 of SNAPC3 project towards a basic surface of tWHD1 consisting of K260, K316 and R418 (Figure 3E). Phylogenetic analysis suggested a role for tWHD1 in binding to the class III-specific Brf2 and SNAPc assemblies, effectively acting as an adaptor allowing the polymerase to engage these vertebrate-specific factors^46^. Indeed, analysis of the cryo-EM maps observed an improvement in local resolution of the TFIIIB:SNAPc density when fully engaged with the RPC5 region in both full-length and mini SNAPc-containing PICs, consistent with its role in stabilizing the complex (Extended Data Figure 8A). In the elongating complex, RPC5 tWHD1 adopts 2 conformations, one which closely resembles its position in the OC^FL^ and a second projecting towards the density^46^. This additional conformation does not appear compatible with PIC assembly as it induces large-scale steric clashes between WH2 of RPC5 and the SNAPC3 wedge region (Extended Data Figure 8B and C), suggesting that this conformation may have additional roles specific to other promoters or stages of transcription.

As previously reported, PSE recognition in our structure was mediated by both SNAPC3 and SNAPC4^36,37^. SNAPC3 presents a basic channel interface which interacts extensively through 3 motifs which wrap around the minor groove from -62 to -56. The DNA interface was formed through 3 conserved motifs consisting of residues 144-150, 190-198 and 348-353 (Extended Data Figure 8D-H). A significant proportion of these interactions were hydrogen bonds between the residues and DNA backbone, however, W350 was inserted into the DNA helix at A -57 in an equivalent position to that observed in structures with the U6-1 PSE sequence (Extended Data Figure 8H)^36^. SNAPC4 contributes to PSE recognition via binding through the conserved MyB DNA binding domain. The MyB domain consists of three alpha helices which mediate binding to the DNA major groove and are often arranged in either 2 or 3 imperfect tandem repeats. SNAPc contains an atypical MyB arrangement, with a single half repeat consisting of 2 helices (R_h_) followed by four canonical MyB repeats (R_a_, R_b_, R_c_ and R_d_)^47^. Previously, R_b_, R_c_ and R_d_ have been identified as important in SNAPC4-mediated PSE recognition^18,47^, however due to flexibility in this region we were able to resolve only the first 2 ⍺-helices of the R_b_ domain in our OC^FL^ structure, which made contact with the DNA phosphate backbone, consistent with previous reports (Figure 3D)^36,37^.

### Sulfo-SDA crosslinking mass spectrometry localises SNAPC5 and SNAPC2 in SNAPc

Inspection of the PIC structure observed extensive flexibility within the SNAPc structure, with large regions not visible in the EM map. To obtain structural information for these absent regions, the DNA-bound SNAPc complex was subjected to sulfo-SDA crosslinking coupled mass spectrometry. The acquired SNAPc-DNA dataset contained 268 crosslinks of which 8 were homomultimeric and 81 heteromeric with a 5% false discovery rate (FDR)(Figure 4A). Mapping of identified crosslinks onto the SNAPc structure found within the OC^FL^ visualized 52 crosslinks, of which 49 were satisfied with crosslinking partners located within 30Å, giving a violation rate of 5.7% (Figure 4B; Extended Data Figure 9A-C). The observation that many of the crosslinks of the DNA-bound SNAPc were consistent with the structure of SNAPc engaged in the PIC suggests that the SNAPc core undergoes very few structural changes upon subsequent TFIIIB and Pol III recruitment and therefore is in an assembly competent conformation upon DNA binding.

**Figure 4.**
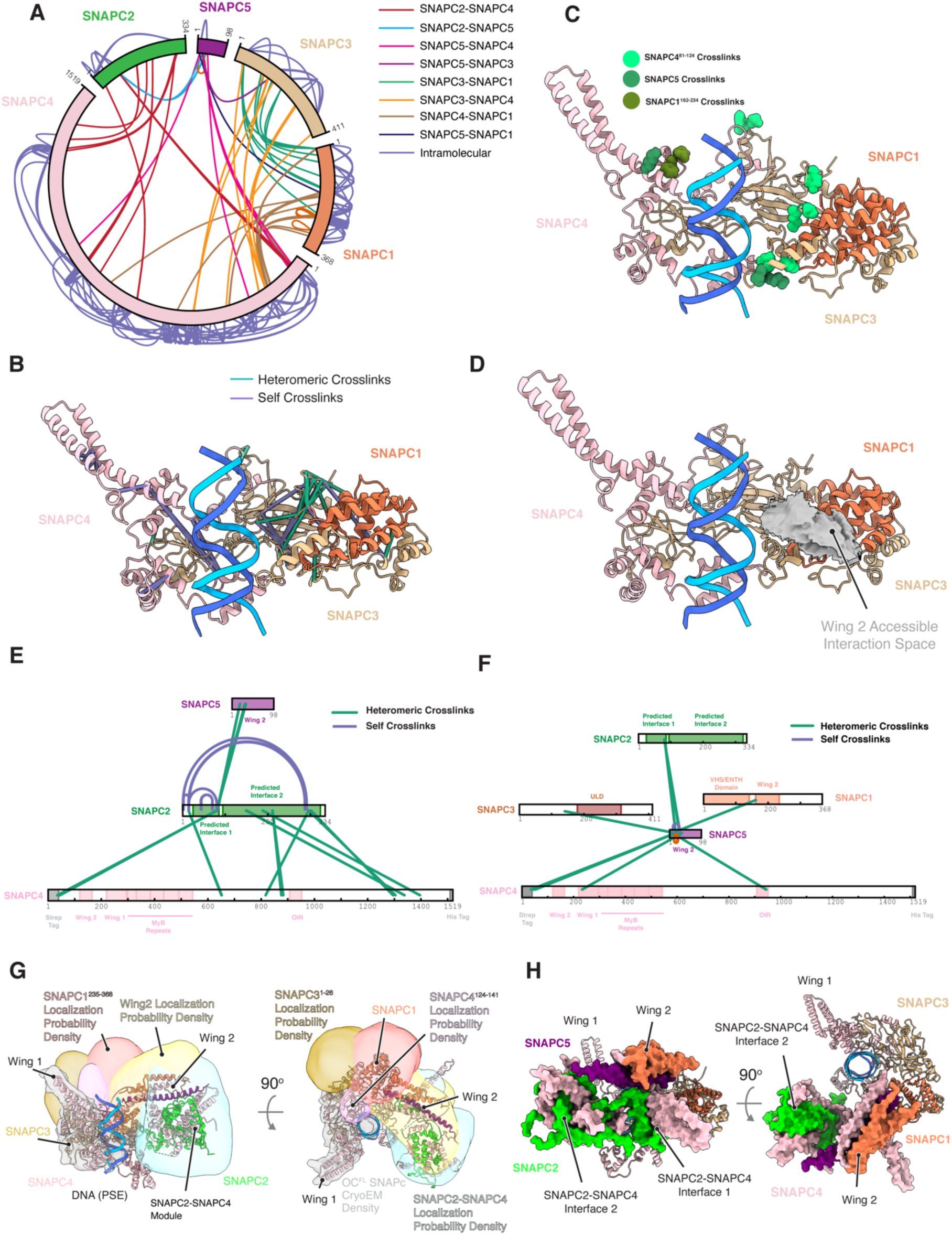
Sulfo-SDA cross-linking mass spectrometric analysis of the SNAPc-DNA complex. (a) Crosslinking mass spectrometry identified 260 crosslinks between proximal residues. Intermolecular crosslinks are shown in the interior, coloured according to subunit interface. Intramolecular crosslinks within each subunit are shown on the exterior. (b) Crosslinks mapped to the SNAPc structure from OC^FL^. (c) SDA half-links with wing 2 are highlighted on the resolved region of SNAPc, coloured according to the cross-linked wing 2 subunit. (d) Visualisation of the crosslink-driven accessible interaction space for wing 2 mapped onto the OC^FL^ SNAPc structure. (e) Crosslinks observed between SNAPC2 and other subunits in SNAPc. Self-crosslinks for SNAPC2 are shown in lilac with heteromeric crosslinks shown in green. (f) Crosslinks observed between SNAPC5 and other subunits in SNAPc. Self-crosslinks for SNAPC5 are shown in lilac with heteromeric crosslinks shown in green. (g) The integrative structural model determined for SNAPc^FL^ at 27Å precision using the sulfo-SDA crosslinks to model the position of wing 2 and the SNAPC2-SNAPC4 regions. For each subunit the localisation probability density derived from the ensemble of models within the dominant cluster is shown with the representative centroid structure fitted within it. (h) General features of the cluster centroid model representation of full length SNAPc structure. Built regions present in the OC^FL^ structure are shown in ribbon, with IMP-modelled regions are shown in surface representation.

In addition, crosslinks were also identified for several regions not directly observed in the PIC model, including the SNAPC2 and SNAPC5 subunits. Previously, SNAPC5 was observed to assemble into a helical bundle termed wing 2^38^, which is not observed in either DNA-bound SNAPc or Pol III PIC structures^36,37^. Several crosslinks were present in the dataset which were consistent with the formation of a wing 2 structure in the Pol III PIC. Multiple crosslinks were identified between SNAPC5, SNAPC1^162–234^ and SNAPC4^81–125^ which were satisfied in the Pol II PIC wing 2 structure, suggesting this region is formed in the absence of additional binding partners and forms a rigid body moving with respect to the rest of SNAPc (Extended Data Figure 9D). Indeed, mapping of wing 2 region crosslinks with the rest of the SNAPc structure and DisVis analysis suggest that this region is highly mobile with multiple crosslinks observed centered over the SNAPC1 subunit (Figure 4C-D). This position in the PIC structure places wing 2 in proximity to the Bdp1 N-terminal region in the PIC structure, which has been implicated in forming direct contacts with SNAPc^29^, allowing for the hypothesis that Bdp1 binds to wing 2 in a manner analogous to TFIIB in the Pol II PIC^38^, with the disordered nature of Bdp1 preventing resolution of this region in Pol III PIC maps. SNAPC2, in contrast, was found to form multiple crosslinks with the flexible C-terminal region of SNAPC4 between residues 651-652, 817-883 and 1295-1396 (Figure 4E). This is consistent with previous mutational studies which localized the SNAPC2 binding site in this C-terminal region^18,20^. However, additional SNAPC4 regions were found to crosslink SNAPC2, including the N-terminal region (encompassing residues 2 to 4). All these SNAPC4 regions crosslinked to SNAPC2 also crosslinked extensively with each other in the SNAPC4 sequence (Extended Data Figure 9E), suggesting that they form a mobile structural module together with SNAPC2. Indeed, alphapulldown analysis yielded a high confidence binding prediction between SNAPC2 and SNAPC4 comprising of 2 distinct interfaces formed between SNAPC2^25–85^-SNAPC4^668–760^ and SNAPC2^95–323^-SNAPC4^1270–1401^, consistent with the observed crosslinks and suggesting the presence of such mobile module (Extended Data Figure 9F-I). Additionally, SNAPC2 crosslinked to the wing 2 subunit SNAPC5 (Figure 4G), which also crosslinked to an N-terminal patch of SNAPC4, suggesting that the SNAPC2-SNAPC4 body could also interact with wing 2.

To further visualize the relative arrangement of these subunits, we used the crosslinking-MS as restraints to derive a model of the relative positioning of these regions in the SNAPc complex using the integrative modelling platform (IMP)^48^. A coarse-grained representation of the complex was determined utilising the determined structure of the SNAPc from OC^FL^, including the MyB binding domains determined for SNAPc^mini^ in complex with the U1 PSE^36^, wing 2 structure from the Pol II PIC (PDB:7zx8)^38^ and the alphafold multimer model of the SNAPC2-SNAPC4 module (Extended Data Table 2). Wing 2, SNAPC2-SNAPC4 interface 1 and interface 2 were defined as rigid bodies which were fully flexible with respect to the SNAPc ’core’ from OC^FL^ . 8146 models were generated which satisfied the restraints, which following clustering identified a single ensemble cluster, representing 70% of the total models (Extended Data Figure 9J-M). The centroid model of this cluster, with a sampling precision of 27Å, was used to interrogate the overall relative positioning of the rigid bodies. Consistent with the DisVis analysis, IMP simulation placed the wing 2 projecting from the N-terminus of SNAPC1 in the SNAPc core, which also contacts the SNAPC2-SNAPC4 module forming a large projection, supporting the suggestion of a large mobile module formed between these regions (Figures 4G-H). Further inspection identified a number of violated crosslinks in the centroid model between wing 2 and the SNAPC2-4 module and the SNAPc ‘core’ (Extended Data Figure 9N), likely due to the high flexibility in this region. Nevertheless, the modelled position of both wing 2 and the SNAPC2-4 module places them on the upstream face of the PIC complex, in proximity to the bound DNA, consistent with the role of these regions in regulating DNA binding affinity and mediating interactions with Oct1 which facilitate Pol III transcription^20,49^.

### SNAPC4 wing 1 forms the major interaction interface with TFIIIB

SNAPc has been characterised as essential in recruiting the class III promoter specific Brf2-TBP-Bdp1 (TFIIIB) assembly to allow for PIC formation and transcription initiation^25–29^. Resolution of the PIC structure allows for a detailed investigation of the mechanism of recruitment. In the OC^FL^ structure, Brf2 is observed to mediate the major interaction with SNAPc via the N-terminal SNAPC4 wing 1 (residues 185-259). Specifically, a small, hydrophobic ⍺-helix (residues 280-288) in the Brf2 C-terminal cyclin repeat projects several hydrophobic residues (A281, W282, V285 and L286) which contact L201, L205, L209, L212 and I242 of SNAPC4 wing 1 (Figure 5A). These SNAPC4 residues form a hydrophobic patch on the surface of SNAPC4 wing 1 which defines the Brf2 binding site (Figure 5A-B). The cyclin ⍺-helix is positioned for SNAPC4 interaction via several stabilizing hydrophobic interactions with the semi-circular Brf2 arch ⍺-helix (residues 291-314). Numerous arch residues (V293, L300, L307) directly contact L280, L283 and L288 in the cyclin ⍺-helix, fixing its positioning to facilitate interaction with SNAPC4 (Figure 5C). Therefore, defining the cyclin ⍺-helix-arch region of Brf2 as the major interface which recruits TFIIIB to SNAPc-bound class III promoters. This is consistent with previous observations, where TBP-binding deficient Brf2 mutants could still form TFIIIB-SNAPc complexes^34^, with specific deletion of the arch helix required to abrogate this complex formation *in vitro*^5^. Indeed, point mutations in the SNAPC4 N-terminus have been recently identified as associated with neurodevelopmental disorders^31^. Mapping of these SNAPC4 patient mutations (D199N and K185E) onto the structure revealed a likely disruption of both the wing 1 structure and its binding to DNA (Figure 5D), highlighting the importance of this region for normal transcription.

**Figure 5.**
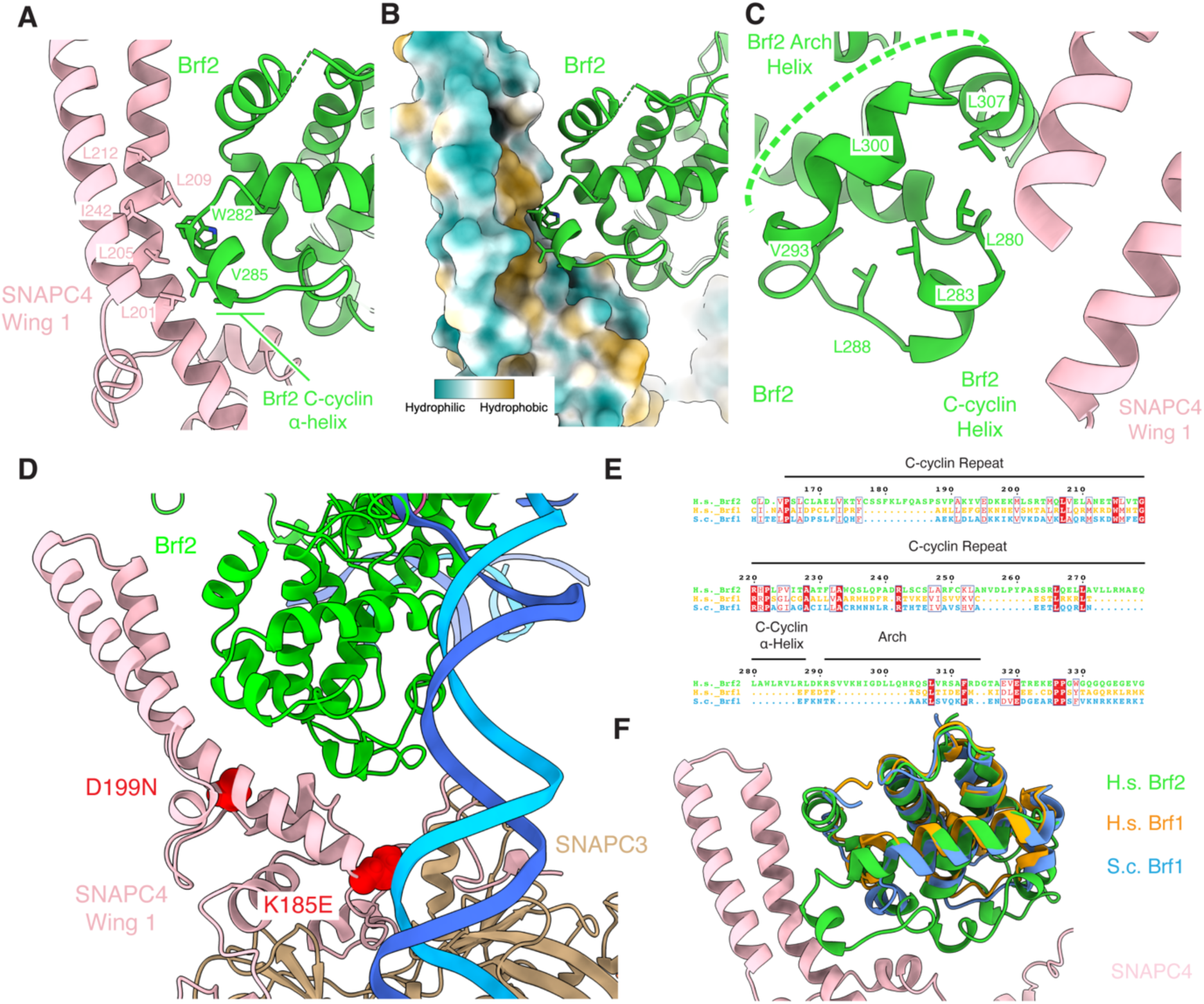
Structural characterization of the TFIIIB-SNAPc interface. (a) The Brf2 C-cyclin-SNAPC4 wing 1 interaction site with interacting residues highlighted. (b) The Brf2-SNAPC4 interface visualised with a hydrophobic surface representation for the SNAPC4 wing 1 region. (c) Visualisation of stabilising interactions between the arch and C-cyclin alpha helix in the Brf2 SNAPC4 interaction module with interacting residues highlighted. (d) Mapping of SNAPC4 neurodevelopmental disorder-associated point mutations in the class III PIC context. (e) Sequence alignment of human Brf2 with human and yeast Brf1, showing the specific presence of the arch and C-cyclin alpha helix in Brf2. Conserved residues are highlighted in red, with sequence similarity labelled with a blue box. (f) Structural alignment of human Brf2 (green) with human (orange) and yeast (blue) Brf1 at the interface with SNAPC4 wing 1 in the human PIC.

Both Brf1 and Brf2 share high homology in the conserved N-terminal Zn-ribbon and cyclin domains with the major differences found in the C-terminal domain (CTD) of both proteins^5^. Sequence comparison of human Brf2 and Brf1 revealed that the Brf2 cyclin ⍺-helix-arch region is absent in Brf1 (Figure 5E). Furthermore, structural alignment between Brf2, the yeast Brf1 structure and the alphafold human Brf1 structure prediction revealed that despite high similarity in the fold observed in the interacting region (Extended Data Table 1), the cyclin ⍺-helix-arch projection required to interact with SNAPC4 wing 1 was specific to Brf2 (Figure 5F), thus ensuring specific recruitment of Brf2-containing TFIIIB to SNAPc-bound promoters.

### The SNAPC4 wing 1 forms a double-sided interaction module which facilitates engagement with both Pol II and Pol III PICs

SNAPc is unique amongst the multi-subunit basal transcription factor apparatus in its ability to engage both Pol II and Pol III at their respective PSE-containing promoters^34,38^. Each polymerase utilizes a distinct set of basal transcription factors which assemble with TBP at the promoter in a mutually exclusive fashion^34^. At Pol III promoters, assembly of the Pol III-specific factors Brf2 and Bdp1 with TBP allows for the formation of the TFIIIB complex which binds SNAPc^5,27–29,34^. However, in the case of Pol II promoters, TBP forms a complex with both TFIIB and TFIIA which mediate both SNAPc and Pol II binding^33,38^. The mutually exclusive nature of TFIIIB or TFIIB-TFIIA assembly underlies the high recruitment specificity observed for these factors across both Pol II and Pol III loci^34^. However, this requires SNAPc to engage with compositionally diverse transcription factor complexes to facilitate recruitment and transcription.

Comparison of the OC^FL^ structure to the SNAPc-containing Pol II PIC structures on U1 and U5 scaffolds revealed a dramatic conformational change with respect to the Pol III complex. Alignment of the structures displayed a 180° rotation of the SNAPc complex around the axis of the promoter DNA (Figure 6A). The principal binding SNAPc site is now formed between a basic patch on the opposing surface of SNAPC4 wing 1 and TFIIA (Figure 6B-C). As TFIIA presents this binding site on the opposing side of the promoter to Brf2, this requires SNAPc to assume a flipped conformation with respect to OC^FL^ to engage the Pol II machinery. In this context, SNAPC4 wing 1 presents a ‘double-sided’ binding motif (Figure 6D), presenting a hydrophobic patch on one side to facilitate Brf2 interaction (Figure 6B) and a basic motif on the opposing face for TFIIA-TFIIB complexes (Figure 6C), giving rise to the observed rotational dispersion. Furthermore, the different nature of binding also prevents non-productive SNAPc engagement with either basal transcription factor complex which would position the SNAPc-PSE binding site away from the promoter. The conformational specificity of SNAPc in both the Pol II and Pol III context is also ensured by steric clashes induced by inappropriate flipping of SNAPc. Alignment of the OC^FL^ to the Pol II PIC structure revealed that in the Pol II conformation, both SNAPC1 (residues 163-171 and 222-232) and SNAPC5 (residues 4-17) in wing 2 would clash with the arch and hydrophobic helix motifs projected by Brf2 in the OC^FL^, with additional clashes observed with the promoter DNA (Figure 6E), thus rendering each mirrored SNAPc conformation compatible with a unique set of transcription factors and defining a ‘Pol II’ and ‘Pol III’ arrangement for SNAPc in their respective PICs.

**Figure 6.**
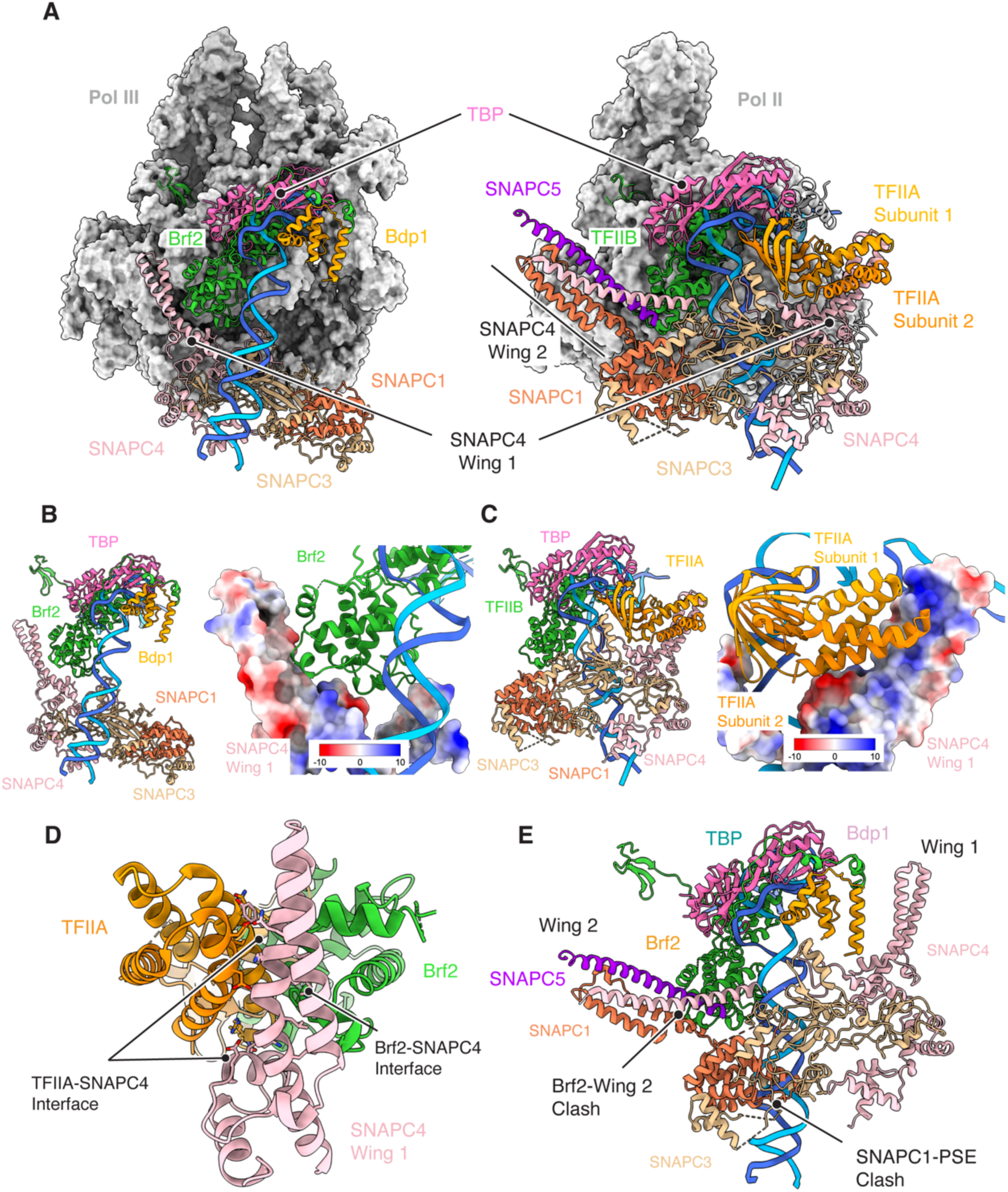
Comparison of the human SNAPc-containing Pol III and Pol II PICs. (a) Visualisation of the human Pol III Pol II PICs with SNAPc and TFIIIB, TFIIB and TFIIA subunits shown in ribbon representation. In both, common SNAPc structural units are highlighted to show the mirror conformations. Pol III subunit surfaces are shown coloured in grey. (b) A focused view of the TFIIIB-SNAPC4 interface relative to the promoter DNA shown in ribbon (*le$*) with a zoomed view of the Brf2-SNAPC4 interface visualized with a colombic surface representation for SNAPC4 wing 1 (*right*). (c) A focused view of the TFIIA-SNAPc interface relative to the promoter DNA shown in ribbon (*le$*) with a zoomed view of the TFIIA-SNAPC4 interface (PDB: 7zx8) visualized with a colombic surface representation for SNAPC4 wing 1 (*right*). (d) Structural alignment of SNAPC4 from Pol III and Pol II PICs, shown are the two distinct interfaces utilized by Brf2 and TFIIA to interact with SNAPC4 wing 1. Interacting residues are highlighted. (e) Steric clashes induced through rotation of SNAPc to adopt the same orientation as that in the Pol II PIC (PDB: 7zx8) in the Pol III PIC context. Clashes between Brf2-wing 2 and SNAPC3-promoter DNA are highlighted.

## Discussion

Here we report the cryo-EM structures of the gene external class III promoter in the OC and MC states assembled with both mini and full length SNAPc transcription factor complexes. We report structures consistent with previous studies of both DNA-bound miniSNAPc and PIC complexes^36,37^. Comparison of OC^FL^ with yeast pol III OC-PIC complexes unveiled unexpected structural differences with the human complex, revealing potential mechanistic differences in transcriptional initiation between the species. Bdp1, the RPC1 rudder and the RPC2 switch III loop displayed significant structural divergence from the yeast counterpart upon transition between apo, PIC and elongating states (Figure 2). In particular, the process of DNA melting and the formation of a full transcription bubble, which in yeast occurs through a series of TFIIIB-dependent allosteric changes of RNA Pol III subunits, appears to be more simplified in the human system and does not require TFIIIB-activation (Figure 2).

The reported structures provide the structural basis for the function of the RPC5 C-terminus (RPC5EXT), which consists of two tandem winged helix domains (tWHD) connected by a large flexible linker^5^. Previously, phylogenetic analysis had suggested that this region had co-evolved with several class III PIC components, effectively functioning as an adaptor which allowed for stable recruitment of the polymerase^46^. Our structures are consistent with this hypothesis, with several charge complementary interfaces identified between RPC5 and SNAPc and full engagement of RPC5EXT by SNAPc required for stabilization of this region (Figure 3D; Extended Data Figure 8A). Whilst this identifies a function for tWHD1, it remains unclear for the tWHD2, which is highly flexible and not present in cryo-EM reconstructions^50^. This tandem domain also co-evolved with the class III machinery and was associated with regulating the stability of the polymerase assembly^46,50^, therefore could potentially recruit additional regulatory proteins to the PIC.

Significantly, structural comparison of our PIC structures with previously reported SNAPc-containing Pol II human PIC revealed a flipping of SNAPc (Figure 6A)^38^. In both cases the wing 1 coiled-coil structure, formed by SNAPC4, was crucial in mediating the interaction by presenting a double-sided interaction motif, with opposing hydrophobic and electrostatic faces able to mediate interactions with Brf2 and TFIIA, respectively (Figure 6B and C). In this way, wing 1 can accommodate the diverse upstream transcription factors recruited by each gene in 2 rotated arrangements directed by the wing 1 face engaged, which also prevents inappropriate, non-productive engagement of the complex. This provides a mechanistic rationalization of how SNAPc can efficiently direct transcription from both the Pol II and Pol III transcriptional apparatus. Hou *et al* have previously reported this flipped orientation of SNAPc between the Pol II and Pol III PICs and proposed a model where the 3 nucleotide (nt) difference in spacing between the PSE and TATA box-TFIIB binding site in Pol III and Pol II promoters causes the relative rotation of SNAPc^37^. However, Pol II snRNA promoters lack a strong TATA box signal for TFIIB-TFIIA positioning, with the selection of the TSS site and the binding of TBP determined by the PSE-bound SNAPc complex which acts as a ‘molecular ruler’ positioning the TFIIB-TFIIA complex at a set distance from the PSE^35,36,38^. As a result, the observed Pol II PIC structure is likely a result of a direct protein-protein interaction formed during SNAPc-mediated recruitment to the promoter, with limited effect from the DNA binding site. Furthermore, the 3 nt difference in promoter spacing described is within the natural variation observed across all SNAPc-containing Pol III genes, with a range of PSE-TATA box spacings between 15 and 18 nt observed and still competent for Pol III transcription (Extended Data Figure 10). Indeed, in vitro transcriptional assays have demonstrated that the U6 promoter can tolerate deletions and insertions of 2 and 3nt respectively between the PSE and TATA box whilst still supporting SNAPc-TFIIIB engagement and Pol III transcription^51^. Furthermore, several Pol III loci (7SK, U6, and RPPH1) have also been observed to be transcribed by Pol II despite their defined Pol III PSE-TATA box spacing^7,34,52^. As a result, the small spacing difference observed between U1 and U6 promoters is unlikely to drive the flipping of the SNAPc. We propose that the flipped orientation is a specific adaptation of SNAPc to engage either with TFIIB and Brf2 complexes at Pol II and Pol III promoters and is governed by a mutually exclusive set of interactions and steric clashes that fix the SNAPc conformation in each complex. Thus, the only determinant of polymerase selectivity is the presence or absence of a TATA box, as previously demonstrated^34^. The presence of a canonical TATA box favors the recruitment of Brf2-TBP complex and, subsequently, RNA Pol III. In the absence of a TATA box, recruitment of TBP-TFIIA prevails, resulting in the subsequent recruitment of RNA Pol II. In both scenarios, SNAPc can interact in a similar manner with the cognate PSE box while adapting two distinct and mutually exclusive conformations to engage with either Brf2-TBP or TFIIA-TBP, resulting in the recruitment of RNA Pol III or Pol II, respectively.

Despite the increasing amount of structural data available for SNAPc and the class III PIC^36–38^, the arrangement of SNAPC2 and SNAPC5 subunits in the SNAPc complex remained elusive. Employing sulfo-SDA crosslink mass spectrometric analysis, the location and arrangement of these subunits was investigated. Combining this crosslinking analysis with alphafold prediction, the SNAPC2 binding site was localized to a structural module in the C-terminus of SNAPC4, consistent with previous mutational analysis which identified the region spanning 1281-1393 of SNAPC4 as the interaction site^18,20^. We therefore confirm this as the binding site of SNAPC2, whilst defining new contact sites in the N and C-termini (Figure 4E). SNAPC5 was also identified as part of the wing 2 helical bundle despite the absence of a defined wing 2 structure in the Pol III PIC and miniSNAPc-DNA^36^ structures, consistent with the requirement for this region to produce a stable SNAPc complex^36^. Multiple crosslinks were identified between both wing 2 and SNAPC2, suggesting that these subunits together form a large structural module which is highly mobile. IMP modelling placed this structure in proximity to the bound DNA, consistent with the role of these subunits in regulating DNA binding affinity^18^. The observed localisation and flexibility of this module allows us to speculate that SNAPC2-SNAPC4 module may regulate DNA access to the binding site hence modulating the SNAPc-DNA binding affinity. Indeed, the identified mobile module encompassed the Oct1 interacting region (OIR) in SNAPC4^18,49^ which was close to several SNAPC2 crosslinking sites (Extended Data Figure 9F) and has been implicated in alleviating this DNA-binding repression and stabilizing SNAPc at the promoter^9,21^, a function which has also been suggested for SNAPC2^18^ leading to the intriguing possibility that Oct1 may interact with a SNAPC2 ’gate’ to regulate SNAPc DNA-binding. In the class III PIC context, this SNAPC2-containing mobile module faces the upstream region, pointing towards the nucleosome and bound Oct1 found across all class III promoters^6^, potentially facilitating this interaction. As a result, future work will focus on characterizing the interaction between SNAPc-containing PICs and upstream regulatory factors to give greater mechanistic insight into class III promoter transcription and a more complete structural description of the complete class III PIC.

## Methods

### Cloning and Protein Expression

All TFIIIB components were expressed recombinantly in Rosetta (DE3) pLysS *Escherichia coli* cells as described previously^5,29^. Briefly, N-terminally His-tagged TBP residues 159-339 (His-TBP_core_) was expressed in a pET45b(+) vector. Following transformation, cells were grown in terrific broth (TB) at 37°C to an optical density (A_600_) of 0.6 and subsequently induced with 0.5mM Isopropyl β-D-1-thiogalactopyranoside (IPTG) at 20°C for 18 hours. Cells were harvested by centrifugation at 2000 *x g* for 30 mins at 4°C and stored at -80°C prior to purification. Full length, C-terminally tagged Brf2 was expressed in a pOPINE construct. Transformed cells were grown in TB at 37°C to a final A_600_ of 0.6 and induced for 4 hours at 30°C with 1mM IPTG. Cells were collected by centrifugation at 2000 *x g* for 20 mins at 4°C and stored at -80°C. C-terminally His-tagged Bdp1 (1-484) was expressed in a pOPINE vector. Cells were grown in TB at 37°C until A_600_ of 0.6. The temperature was reduced to 16°C and cultures subsequently induced with 0.5mM IPTG once A_600_ reached 0.8. Induction was carried out for 16 hours and cells collected by centrifugation at 2000 *x g* for 20 mins at 4°C. Cell pellets were stored at -80°C.

Full length SNAPc (SNAPc^FL^) and mini SNAPc (SNAPc^mini^) were expressed as previously using the bIGBac baculovirus expression system^38^. Briefly, for SNAPc^FL^ expression, a double-tagged SNAPC4 subunit, carrying a TEV-cleavable N-terminal strep and C-terminal His tag, SNAPC1, SNAPC2, SNAPC3 and SNAPC5 were each sub-cloned into a pLIB vector and subsequently combined into a pBIG2ab expression construct by Gibson Assembly (NEB) using the biGBac expression system. For SNAPc^mini^ construction, an N-terminal TEV-cleavable Strep and His tag were added to SNAPC4 (1-505aa) and SNAPC3 respectively. These constructs were sub-cloned into pLIB vectors alongside SNAPC1 (1-268aa), SNAPC2 and SNAPC5 and subsequently combined into a final pBIG2ab construct for expression. Following cloning, both SNAPc^FL^ and SNAPc^mini^ pBIG2ab constructs were used for bacmid production via transformation into DH10 EMBacY cells. For baculovirus generation, bacmid was isolated and transfected into 2ml of adherent Sf9 cells (Thermo Fisher Scientific) at 0.5x10^6^ cells/ml density in Insect-XPRESS media (Lonza) in a 6-well plate. Cells were incubated at 27°C for 72 hours and supernatant (total volume of 2ml), containing the baculovirus, was collected producing the P1 virus. Viral amplification was achieved by P1 infection of a 25ml culture of Sf9 cells growing in suspension at a cell density of 0.5x10^6^ cells/ml in Insect-XPRESS media (Lonza). Cells were incubated at 27°C at 130 r.p.m. for 5 days and supernatant collected following centrifugation of the cultures at 1000 *x g* for 15 mins at 4°C. This produced the P2 virus which was stored at 4°C. Large-scale protein expression was carried out through addition of 2ml of P2 viral stock to 500ml of High Five cells grown to 0.5x10^6^ cells/ml in Insect-XPRESS media (Lonza) in a roller bottle flask (Greiner). Cells were grown at 27°C for 3 days at 130 r.p.m. and harvested through centrifugation at 1000 *x g* for 20 mins at 4°C. Cell pellets were resuspended in phosphate buffered saline (PBS), centrifuged as previously and the resulting washed pellet stored at -80°C prior to purification.

Tagged endogenous human RNA polymerase III was purified from HEK293T cells CRISPR/Cas9 modified to include a C-terminal mCherry-Twin Strep-His-P2A tag on the RPAC1 (POLR1C) subunit, as described previously^46^. To produce sufficient material for purification, large-scale cell cultivation was carried out in a 10L Autoclavable Bioreactor (ez2-Control Bioreactor, Applikon) using a batch cultivation in Dynamis AGT Medium (Gibco) supplemented with 1% penicillin-streptomycin, 4mM stable glutamine (Euroclone), 0.3% Pluronic F-68 non-ionic surfactant (Gibco), and 90ppm silicone-based Antifoam C Emulsion (Sigma-Aldrich). Cultivation conditions included a temperature of 37°C, pH 7.0 (controlled by CO_2_), and dissolved oxygen at 40%. The latter was controlled by sparger gassing with airflow ranging from 100 to 500 ml/min, along with a second cascade of O_2_ gassing from 0 to 100 ml/min. Agitation was maintained at 100 r.p.m. using a three-pitched blade impeller. Initially, cells underwent subculture and expansion until reaching a density of 6–8 × 10^6^ viable cells/ml. Subsequently, the cells were used to inoculate the bioreactor at a concentration of 0.5-1 × 10^6^ viable cells/ml, depending on the desired cultivation time. Harvesting took place at 8–10 × 10^6^ viable cells/ml.

### Protein Purification

For purification, TBP_core_ pellets were resuspended in lysis buffer (50 mM HEPES pH 8.0, 500 mM NaCl, 10 mM Imidazole, 5mM β-mercaptoethanol, 1mM MgCl_2_ supplemented with 1:500 protease inhibitor cocktail set III (Calbiochem) and DNAseI (20mg/ml)). Lysis was carried out via sonication at 40% amplitude for a total of 15 mins on ice and the resulting lysate cleared via centrifugation at 48000 *x g* for 1 hour. The collected supernatant was further cleared via successive filtration using a 12µm and then a 0.45µm filter. The lysate was loaded onto a 5ml HisTrap HP (Cytiva) pre-equilbrated with Buffer A1 (50 mM HEPES pH 8.0, 500mM NaCl, 10% (v/v) glycerol, 5 mM β-mercaptoethanol, 10 mM Imidazole) and washed with 12% (v/v) of Buffer B (50 mM HEPES pH 8.0, 500mM NaCl, 10% (v/v) glycerol, 300 mM Imidazole, 5 mM β-mercaptoethanol). TBP_core_ was then eluted with in 100% buffer B and the elute loaded directly on a 5ml Heparin HiTrap HP column (Cytiva) pre-equilibrated with a 75%:25% (v/v) mixture of Buffer H1 (50mM HEPES pH 8.0, 1mM DTT, 10% (v/v) glycerol) and Buffer H2 (50mM HEPES pH 8.0, 2M NaCl, 1mM DTT, 10% (v/v) glycerol). The protein was eluted via a linear gradient from 25% to 75% (v/v) Buffer H2. Eluted protein was supplemented with 1mM MgCl_2_ and incubated overnight at 4°C with 300µg His-3C protease per 7.5mg purified TBP_core_. Following overnight incubation, the mixture was applied to a 5ml HisTrap HP column (Cytiva) pre-equilibrated with Buffer A2 (50 mM HEPES pH 8.0, 500mM NaCl, 10% (v/v) glycerol, 5 mM β-mercaptoethanol, 30 mM Imidazole) and the flow-through collected. Purified TBP_core_ was then concentrated using a Vivaspain6 (10k MWCO, Sartorius) and stored at -80°C.

Both Full length Brf2-His and Bdp1(1-484)-His were purified following the same protocol, as previously described^5,29^. Initially, pellets were resuspended in lysis buffer (50 mM HEPES pH 8.0, 750 mM NaCl, 10 mM Imidazole, 5mM β-mercaptoethanol, 1mM MgCl_2_ supplemented with 1:500 protease inhibitor cocktail set III (Calbiochem) and benzonase (Sigma-Aldrich)). Cells were lysed via sonication at 40% amplitude for a total of 15 mins on ice and the resulting lysate cleared via centrifugation at 48000 *x g* for 1 hour and subsequent filtration. The lysate was loaded on a 5ml HisTrap HP column (Cytiva) pre-equilibrated with lysis buffer and then washed with 83%:17% (v/v) mixture of lysis buffer: buffer B1 (50 mM HEPES pH 8.0, 750mM NaCl, 300 mM Imidazole, 5 mM β-mercaptoethanol). His-tagged protein was eluted in buffer B1 and then diluted two-fold in buffer H1 (50mM HEPES pH 8.0, 1mM DTT) to reduce the salt concentration to approximately 375mM NaCl. This sample was loaded on a Heparin HiTrap HP column (Cytiva) pre-equilibrated with a 83%:17% (v/v) mixture of buffer H1:buffer H2 (50mM HEPES pH 8.0, 2M NaCl, 1mM DTT). The protein was eluted in a linear gradient from 17% to 75% buffer H2. Protein fractions were pooled, concentrated, and subject to gel filtration chromatography using a Superdex 200 16/600 column pre-equilibrated in 50mM HEPES pH 8.0, 200mM NaCl, 10% (v/v) glycerol and 2mM DTT. Protein fractions were pooled and stored at -80°C.

SNAPc^FL^ pellets were resuspended in buffer A1 (50mM HEPES pH 8.0, 750mM NaCl, 10% (v/v) glycerol, 10mM imidazole, 7.5mMβ-mercaptoethanol, 2mM MgCl_2_) supplemented with 1:500 protease inhibitor cocktail set III (Calbiochem) and benzonase (Sigma-Aldrich). Pellets were homogenized on ice using a dounce homogenizer and lysed by sonication for 5 min at 40% amplitude. Whole cell lysate was cleared via centrifugation at 48 000 *x g* and filtration. Cleared lysate was loaded onto a 5ml HisTrap HP (Cytiva) pre-equilibrated with lysis buffer. Protein was washed initially with buffer B1 (50mM HEPES pH 8.0, 500mM NaCl, 10% (v/v) glycerol, 50mM imidazole, 7.5mM β-mercaptoethanol, 10mM O-phospho-L-serine) and then buffer B2 (50mM HEPES pH 8.0, 1.25M NaCl, 10% (v/v) glycerol, 50mM imidazole, 10mM β-mercaptoethanol, 10mM O-phospho-L-serine) followed by a final wash with buffer B1. Protein was then eluted in His-elution buffer (50mM HEPES pH 8.0, 500mM NaCl, 10% (v/v) glycerol, 300mM imidazole, 10mM β-mercaptoethanol). The protein was subsequently diluted with buffer HA (50mM HEPES pH 8.0, 10% (v/v) glycerol, 1mM DTT) to a final salt concentration of 250mM. This sample was loaded onto a 5ml Heparin HiTrap HP column (Cytiva) pre-equilibrated with a 87.5%:12.5% (v/v) mixture of buffer HA:buffer HB (50mM HEPES pH 8.0, 2M NaCl, 10% (v/v) glycerol, 1mM DTT) and then eluted in a linear gradient from 20% -60% buffer HB. Eluted protein was pooled, concentrated and loaded onto a Superose6 Increase 10/300GL (Cytiva) pre-equilibrated in 50mM HEPES pH 8.0, 50mM NaCl, 2mM DTT. Recovered protein was concentrated and stored at -80°C.

SNAPc^mini^ pellets, corresponding to 3L of expression culture, were resuspended in 75ml lysis buffer (50mM HEPES pH 8.0, 750mM NaCl, 10% (v/v) glycerol, 7.5mM β-mercaptoethanol, 2mM MgCl_2_ supplemented with 1:500 protease inhibitor cocktail set III (Calbiochem) and DNaseI(Roche)). The cell suspension was homogenized on ice using the dounce homogenizer, following which benzonase (Sigma-Aldrich) was added and incubated for 30 min at 4°C. Lysis was carried out via sonication for a total of 2 min at 40% amplitude and the lysate cleared through centrifugation at 235 000 *x g* for 40 min at 4°C. The partially cleared lysate was collected and passed successively though a 1.2µm and then 0.45µm filter (Merck-Millipore). The fully cleared lysate was then loaded on a 5ml StrepTrap XT column (Cytiva) pre-equilibrated with buffer A1 (50mM HEPES pH 8.0, 500mM NaCl, 10% (v/v) glycerol, 7.5mM β-mercaptoethanol). The column was washed first with buffer A1, then buffer B1 (50mM HEPES pH 8.0, 500mM NaCl, 10% (v/v) glycerol, 7.5mM β-mercaptoethanol, 10mM O-phospho-L-serine) followed by a final wash in buffer A1. SNAPc^mini^ was eluted in strep elution buffer (50mM HEPES pH 8.0, 500mM NaCl, 10% (v/v) glycerol, 7.5mMβ-mercaptoethanol, 50mM Biotin). The resulting elute was diluted two-fold with buffer HA (50mM HEPES pH 8.0, 10% (v/v) glycerol, 1mM DTT) and loaded onto a 5ml Heparin HiTrap HP column (Cytiva) pre-equilibrated in 50mM HEPES pH 8.0, 250mM NaCl, 10% (v/v) glycerol, 1mM DTT and subsequently eluted with buffer HB (50mM HEPES pH 8.0, 2M NaCl, 10% (v/v) glycerol, 1mM DTT). The SNAPC3 His tag was cleaved via overnight incubation at 4°C with 500µg TEV protease per 10mg of SNAPc^mini^. Following cleavage, the SNAPc^mini^ sample was recovered and supplemented with 30mM imidazole. This was then loaded on a 5ml HisTrap HP (Cytiva) pre-equilibrated with buffer A and eluted with a 0-100% linear gradient with buffer B. The flow-through was collected, concentrated, and loaded on a HiLoad Superdex 200 16/600pg (Cytiva) pre-equilibrated with 50mM HEPES pH 8.0, 250mM NaCl, 10% (v/v) glycerol, 2mM DTT. Protein fractions were pooled, concentrated and stored at -80°C.

Pellets corresponding to 5L of CRISPR/Cas9 modified HEK293T cells with tagged POLR1C-mCherry-Twin Strep-His-P2A were used for endogenous human RNA Pol III purification. Pellets were resuspended in 300ml of lysis buffer (25 mM HEPES pH 8.0, 150 mM (NH_4_)_2_SO_4_, 5% (v/v) glycerol, 2 mM β-mercaptoethanol, 5 mM MgCl_2_, 10µM ZnCl_2_, 20mM Imidazole) supplemented with 1:500 protease inhibitor cocktail set III (Calbiochem) and bezonase (Sigma-Aldrich). Lysis was carried out via passage through a dounce homogenizer on ice and sonication for 5 min at 40% amplitude. The resulting lysate was cleared via centrifugation at 235 000 *x g* using a Ti-45 rotor at 4°C. Cleared lysate was collected and filtered prior to loading on a 5ml HisTrap Excel column (Cytiva) pre-equilibrated with Ni-buffer A (25 mM HEPES pH 8.0, 150 mM (NH_4_)_2_SO_4_, 5% (v/v) glycerol, 2 mM β-mercaptoethanol, 5 mM MgCl_2_, 20mM imidazole). The sample was eluted with Ni-Buffer B (25 mM HEPES pH 8.0, 100 mM (NH_4_)_2_SO_4_, 5% (v/v) glycerol, 2 mM β-mercaptoethanol, 5 mM MgCl_2_, 300mM Imidazole) and immediately loaded on a 1ml ResourceQ column (Cytiva) pre-equilibrated with RQ-buffer A (25 mM HEPES pH 8.0, 100 mM (NH_4_)_2_SO_4_, 5% (v/v) glycerol, 10 mM DTT, 5 mM MgCl_2_). Pol III was eluted in a linear gradient from 0% to 100% RQ-buffer B (25 mM HEPES pH 8.0, 1M (NH_4_)_2_SO_4_, 5% (v/v) glycerol, 10 mM DTT, 5 mM MgCl_2_) over 40 column volumes (CVs). Pol III fractions were pooled, concentrated and used either immediately for PIC assembly or stored at -80°C.

### Human Class III PIC Preparation

Both SNAPc^FL^ and SNAPc^mini^ human class III PICs (PIC^FL^ and PIC^mini^) were assembled on the U6-2 scaffold (Non-Template strand: 5’ -AAACC GACCATAAGTTATCCTAACCAAAAGATGATTTGATTGAAGGGCTTAAAATAGGTGTGACAGTAACCCTTGAGTCGTGCTCGCTTCGGCAGCAC -3’; Template strand: 5’ – GTGCTGCCGAAGCGAGCACGACTCAAGGGTTACTGTCACACCTATTTTAAGCCCTTCAATCAAATCATCTTTTGGTTAGGATAACTTATGGTCGGTTT -3’) annealed in 50mM Tris-HCl pH 8.0, 1mM EDTA. The formation of the PIC used all components in excess relative to the human polymerase with assembly carried out in a sequential manner. First, 250pmol of annealed U6-2 DNA scaffold was incubated with 1nmol of either SNAPc^FL^ or SNAPc^mini^. This was incubated for 2 hours on ice (in the case of SNAPc^FL^) or for 15 mins at 25°C (in the case of SNAPc^mini^). Next, 500pmol of TBP_core_ and 500pmol Brf2 were added and incubated at 25°C for a further 15 mins. This was followed by addition of 500pmol of Bdp1(1-484) and an additional incubation step of 15 mins at 25°C. 100pmol of purified human Pol III was subsequently added and incubated for 5 mins at 25°C. The mixture was then diluted twelve-fold with assembly buffer (50mM HEPES pH 8.0, 150mM NaCl, 10mM MgCl_2_, 3mM DTT) to reduce the salt concentration and subsequently concentrated using a Vivaspin6 100k MWCO (Sartorius). The assembly was incubated for a final 30 mins at 25°C. The reconstituted class III PICs were subjected to 15-35% glycerol gradient ultracentrifugation prepared in GraFix buffer (25mM HEPES, 150mM NaCl, 5mM MgCl_2_, 3mM DTT) with simultaneous cross-linking with 0.01% (v/v) glutaraldehyde using GraFix at 197 000 *x g* for 18 hours at 4°C in a SW-Ti 41 rotor. The gradient was fractionated into 200µl aliquots and quenched with 100mM Tris-HCl pH 8.0. Fractions containing the complex, as assayed by SDS-PAGE, were pooled and concentrated using a Vivaspin6 100k MWCO (Sartorius). The resulting sample was used immediately for grid preparation.

### Cryo-EM Sample Preparation and Data Collection

Class III PIC samples were prepared on Quantifoil R 1.2/1.3 300-mesh Cu grids coated in-house with a Leica ACE600 setting a 1.5nm thickness target for deposition of a continuous layer of amorphous carbon. Grids were glow-discharged at 30mA for 60s using a GloQube instrument (Quorum Technologies) prior to sample addition. For PIC^FL^, a final concentration of 0.03% (v/v) Octyl-β-glucoside was added immediately prior to grid preparation. All samples were prepared using the VitroBot Mark IV system (Thermo Scientific). For both PIC^FL^ and PIC^mini^ samples, 3µl was applied to the grid and incubated for 60s at 4°C. Grids were blotted for 3.5s at blot force -5 with a drain time of 0.5s and plunge frozen in liquid ethane.

Data were collected using an FEI Titan Krios G4 transmission electron microscope (Thermo Scientific) operating at 300keV equipped with a Falcon 4i (Thermo Scientific) direct electron detector and EF-TEM mode using a SelectrisX (Thermo Scientific) post-column energy filter operating at a slit width of 10eV. All datasets were collected using EPU automated acquisition software in electron counting mode at a nominal magnification of 130 000x corresponding to a calibrated pixel sampling of 0.955 Å pixel^-1^. All datasets were collected in Electron Event Representation (EER) file format. PIC^FL^ movies were collected with a total dose of 50*e^-^* Å^-2^ over a 5.3s exposure time, giving a dose rate of 8.57 *e^-^* pixel^-1^ s^-1^ which was fractionated over 34 frames yielding a dose of 1 *e^-^* Å^-2^ per frame. 17291 total movies were collected over a defocus range of -0.75µm to -1.5µm. For PIC^mini^, movies were collected with a total dose of 50 *e^-^* Å^-2^ over a 6.26s exposure time, yielding a dose rate of 7.29 *e^-^* pixel^-1^ s^-1^. Movies were fractionated into 38 frames, giving a dose of 1 *e^-^* Å^-2^ per frame. 19060 total movies were collected with a defocus range of -1µm to -2.5µm.

### Cryo-EM Data Processing

Frame alignment and dose weighting was carried out using MotionCor2^53^. CTF estimation was carried out using CTFFIND4.1.4^54^, particle picking and 2D classification was carried out in RELION-4.0 for the PIC^FL^ dataset^55^. Initially, 1 252 298 particles were selected using TOPAZ^56^, producing a final combined particle set of 121 292 following several rounds of 2D classification. These particles were used to produce an *ab initio* reference in RELION-4.0^57^ which was used for 3D classification of the particle set into 3 classes. This yielded a single class of 71 347 particles which corresponded to the PIC^FL^ structure. A consensus refinement yielded a reference which was imported into Cryosparc v4.3.1^58^ together with the particle set for further 3D classification without alignment into 5 classes. This produced 3 classes which corresponded to different states of PIC promoter engagement. Classes 1 (29 968 particles) and 2 (22 559 particles) corresponded to an open complex (OC-PIC^FL^) with a closed (Class 1) and open (Class 2) RPC1 clamp. The third class (18 715 particles) displayed a significantly more extended RPC1 clamp and disordered downstream DNA, so was labelled a melting complex (MC-PIC^FL^) as previously observed by Hou *et al*^37^. To improve the TFIIIB-SNAPc^FL^ density, masked 3D classification without alignment focused around the upstream TFIIIB-SNAPc^FL^ region was carried out into 2 classes for each PIC state. This identified distinct OC-PIC^FL^ complexes representing a fully engaged TFIIIB:SNAPc^FL^ with a closed RPC1 clamp, termed OC1^FL^ (15 661 particles), an open clamp state with a partially engaged TFIIIB:SNAPc termed OC2^FL^ (11 036 particles). The same classification of the MC-PIC^FL^ produced an MC state with a fully engaged TFIIIB-SNAPc^FL^ (9 476 particles). Subsequent homogenous refinement in Cryosparc v4.3.1 yielded reconstructions of 3.26Å (OC1^FL^), 3.40Å (OC2^FL^) and 3.51Å (MC^FL^) resolution. Local resolution estimation and filtering was carried out in Cryosparc v4.3.1^58^.

For the PIC^mini^ dataset, frame alignment, dose weighting and CTF estimation was carried out on-the-fly using the Cryosparc Live functionality in the Cryosparc v4.3.1 software suite^58^. Particle picking, 2D classification, *ab initio* reconstruction and 3D classification were carried out in Cryosparc v4.3.1. Following template-based picking, an initial dataset of 1 014 657 particles were selected, yielding a final particle set of 95 438 following several rounds of 2D classification. This was then subject to 3D classification without alignment into 5 classes using a consensus refined *ab initio* reference generated from the same particle set. Classes 1 and 2 represented two distinct states of the class III OC-PIC^mini^ with an open (Class 1) or closed (Class 2) RPC1 clamp. Both reconstructions displayed poor density for the upstream bound TFIIIB-SNAPc^mini^ complex so was subject to a masked 3D classification without alignment focused on TFIIIB-SNAPc^mini^. The TFIIIB-SNAPc^mini^ density did not improve in the ‘clamp open’ map, however, the same classification with the RPC1 ‘clamp closed’ population produced 2 classes corresponding to a fully engaged TFIIIB-SNAPc^mini^ (Class 1, OC1^mini^) reconstructed from 15 386 particles and partially engaged TFIIIB-SNAPc^mini^ (Class 3, OC2^mini^) reconstructed from 13 256 particles. Each class was exported to RELION-4.0 for refinement, yielding maps of 3.76Å and 4.14Å resolution, respectively. Due to variation in the local resolution of the map, local resolution was estimated and filtered using the functionality in RELION-4.0^55^. In each case, the reconstruction was assessed using the 3DFSC remote processing server^59^.

### Model Building and Refinement

An initial model was built through rigid fitting of the elongating human RNA polymerase III (PDB: 7AE1) structure and the U6-2 promoter bound TFIIIB (consisting of Brf2, Bdp1 (1-484) and TBP_core_) structure (PDB: 5N9G) in UCSF Chimera^5,29,46,60^. For SNAPc, an alphafold prediction for the SNAPc complex was calculated using the multimer mode in Alphafold v2.2.4^61,62^. The resulting high confidence prediction was rigidly fitted into the remaining density in UCSF Chimera^60^. Initially, the upstream DNA was constructed through extension of the DNA from the TFIIIB structure which was then fitted into the corresponding density using COOT.^63^ The register of the fitted downstream DNA was determined using the contacts with the polymerase as a reference. The sequence of this model was mutated in COOT to correspond with the U6-2 scaffold sequence. The initial model was refined using PHENIX real space refinement^64^. At this point, model regions not observed in the map density were deleted and the resulting model subject to a further refinement in PHENIX^64^. Following this initial refinement, models were iteratively refined using both PHENIX and ISOLDE^65^, producing the final models which were assessed using Molprobity^66^.

### Cross-Linking Mass Spectrometry

For sample preparation, 100 µg of SNAPc^FL^ was incubated with DNA for 30min on ice to assemble a protein/DNA complex. It was then mixed with a freshly dissolved sulfo-SDA at a ration of 1:2 (w/w). Reaction was carried out in the dark for 1h at room temperature on shaker to react the NS-ester side of the crosslinker. The diazirine group was then photo-activated using ultra-violet light irradiation for 20 mins. A UVP CL-3000L UV crosslinker at 365 nm was used for UV photoactivation. Reaction was quenched using 50 mM Tris pH 8.0. For peptide generation, in-solution digestion was then carried out using an iST bottom-up proteomic sample preparation kit (PreOmics, Art. Nr. P.O.00027) according to manufacturer’s instructions. The resulting peptides were fractionated using Superdex 30 Increase 3.2/300 column in 30% acetonitrile, 0.1% trifluoroacetic acid.

Approximately 1µg of peptides were injected for each liquid chromatography-mass spectrometry (LC-MS) acquisition. The LC-MS platform consisted of a Vanquish Neo system (ThermoFisher Scientific) connected to an Orbitrap Eclipse Tribrid mass spectrometer (ThermoFisher Scientific) operating under Tune 4.0.491. Mobile phases consisted of 0.1% v/v formic acid in water (mobile phase A) and 0.1% formic acid in 80% acetonitrile/water v/v (mobile phase B). Samples were dissolved into 4% mobile phase B. The samples were separated on an EASY-Spray PepMap Neo column (75 µm x 50 cm) (ThermoFisher Scientific) with a flow of 300 nl/min. Peptides were separated on 110 minute gradients designed to match the hydrophobicity of the various SEC fractions.

MS1 spectra were acquired with a resolution of 120,000 and 50ms maximum injection time. Source RF lens was set to 35%. Dynamic exclusion was set to single count in 45 seconds. The duty cycle was set to 2.5 seconds. A precursor charge filter was set to z=3-7. Precursors were selected based on a data-dependent decision tree strategy^67^prioritizing charge states 4-7 and subjected to stepped HCD fragmentation with normalized collision energies of 20,27,30. MS2 scans were acquired with a normalized gain control target of 250% with maxmum injection time of 250ms and an orbitrap resolution of 60,000. For each fraction, double injections were carried out with the second injection performed with a single count dynamic exclusion of 10 seconds.

Raw files were converted to mgf format using ProteoWizard MSconvert (version 3.0.22314)^68^. A recalibration of the MS1 and MS2 m/z was conducted based on high-confidence (<1% false discovery rate) linear peptide identifications using xiSEARCH (version 1.6.745)^69^. Crosslinking MS database search was performed in xiSEARCH (version 1.7.6.7) on a database comprising SNAPC complex and common mass spec contaminants derived from MaxQuant ^70^searched with 4 missed tryptic cleavages. Precursor mass error tolerance was set to 3ppm and MS2 error tolerance to 5ppm. The search included methionine oxidation, SDA loop link (+82.04186484 Da) as fully variable modification and hydrolized SDA (+100.0524), SDA-Tris (202.107934) as variable modifications on linear peptides. The SDA crosslinker was defined as cleavable^71^. The search was set to account for noncovalent gas-phase associations. Prior to FDR estimation, search results were filtered to remove peptides consecutive in sequence. Results were filtered to 5% FDR at the residue pair level using xiFDR^72^ (version 2.1.5.2) and the “boost” feature to optimize thresholds at the lower error levels was enabled to maximise heteromeric crosslinks. Results were exported in mzIdentML format and uploaded to xiview.org for visualization. Pseudobonds files were downloaded and visualized on the structural model using Chimera X.

### Accessible Interaction Volume Analysis (DisVis)

The accessible interaction volume for Wing 2 in the SNAPc complex was carried out using DisVis^73^. The SNAPc core structure from the OC^FL^ reconstruction was defined as the fixed chain with the wing 2 structure from the Pol II U1-snRNA PIC (PDB:7zx8)^38^ used as the scanning chain. Scanning was carried out with a 1Å grid spacing and rotational sampling interval of 15°. The permitted Cβ-Cβ distance range was set to 2.5-22.5Å.

### Integrative Protein Modelling Platform (IMP)

For IMP modelling, the SNAPc complex from the OC^FL^ structure (consisting of SNAPC1 1-141, SNAPC3 27-411 and SNAPC4 142-383) was extracted together with the corresponding EM density as the starting point for modelling. The SNAPC4 MyB DNA binding domains were obtained from the SNAPcmini-U1 PSE structure (PDB:7xur, residues 385-502)^36^ with the wing 2 structure derived from the Pol II U1-snRNA PIC structure (PDB:7zx8)^38^. Interpretation of the SNAPC2-SNAPC4 interface was carried out using alphapulldown structure prediction in the ‘all verus all’ modality^74^. High confidence models were selected based on iptm scores and visualized in UCSF Chimera^60^. Predicted aligned error (PAE) plots visualized using PAE viewer^75^. The resulting high-confidence region was defined as the double-interface SNAPC4-SNAPC2 module. The models were coarse grained and defined as separate rigid bodies. The SNAPc core from OC^FL^ was defined as a single rigid body with its position fixed by the corresponding EM density during sampling. The MyB domain was placed according to the position defined in the SNAPcmini-PSE DNA bound structure (PDB:7xur) and included in this fixed body^36^. Wing 2 (consisting of SNAPC1 (residues 162-234), SNAPC4 (residues 81-124) and SNAPC5), SNAPC2-SNAPC4 interface 1 (SNAPC2 residues 5-85 and SNAPC4 residues 666-759) and SNAPC2-SNAPC4 interface 2 (SNAPC2 residues 95-323 and SNAPC4 residues 1264-1394) were defined as coarse-grained rigid bodies which moved with respect to the SNAPc ’core’-MyB body. Regions for which no structure could be ascribed were coarse grained as beads and maintained as fully flexible. Rigid body definitions, residue ranges and bead sizes are available in extended data table 2. Subsequently, these structural units, using the crosslinking MS data as restraints were used to build an integrative protein structural model for the full SNAPc complex using the IMP 2.20 suite^48^. A total of 268 crosslinks (81 heteromeric) were used as a distance restraint utilising the Bayesian CrossLinkingMassSpectrometryRestraint, with the scoring function inflection point set at 22Å for Cα-Cα lengths and a weight of 1. The connectivity and excluded volume restraints were set with weights of 2 and 1, respectively. Sampling was performed by Replica Exchange Gibbs sampling in IMP 2.20, using 8 replicas in a temperature range between 1.0 and 5.0 in 4 independent runs with initial configurations randomised in each case. Each sampling step allowed for a 0.1 rad rotation and 6Å translation of each bead and rigid body. A model was saved every 10 sampling steps for a total of 10 000 frames during sampling. The resulting population of models was scored to select for agreement with the sulfo-SDA crosslinking restraints. The population of 320 000 sampled configurations was scored according to the crosslinking MS restraint (<220) and excluded volume restraint (<17) cutoffs, yielding an 8259 model subset. Sampling exhaustiveness and completeness was determined as previously^76^ and the sampling precision assessed by RMSD, followed by clustering. 3 ensemble clusters were identified at 44Å global Cα sampling precision, computed on all regions with enabled movers, with cluster 0 representing the dominant cluster containing 70% of total models (5766 models). The precision of cluster 0 was determined by average RMSD to the cluster centroid as 27Å computed on all regions with enabled movers, including fully coarse-grained regions. The resulting centroid model from the most populated cluster was used for all subsequent analysis.

### Electrophoretic Mobility Shift Assay (EMSA)

For EMSA DNA binding analysis, all oligonucleotide sequences were derived from the U6-2 sequence. For TFIIIB assembly, a Cy5-labelled scaffold consisting of the TATA box and flanking regions (Non-Template: 5′ Cy5-ATTTGATTGAAGGGCTTAAAATAGGTGTGACAGTAACC 3′; Template: 5′ GGTTACTGTCACACCTATTTTAAGCCCTTCAATCAAAT 3′) were annealed in 10mM Tris-HCl pH 7.5, 0.5mM EDTA and used directly for binding in a 20µl reaction volume. 1pmol of annealed construct was mixed with 5 pmol TBP_core_ and 7pmol Brf2. To this, Bdp1 (1-484) was titrated in the range of 2-8pmol for full TFIIIB assembly. Complexes were incubated at 25°C for 1 hour. SNAPc DNA binding was assessed using a Cy5-labelled U6-2 DNA sequence annealed as previously (Non-Template: 5’ Cy5-AAACC GACCATAAGTTATCCTAACCAAAAGATGATTTGATTGAAGGGCTTAAAATAGGTGTGACAGTAACCCTTGAGTCGTGCTCGCTTCGGCAGCAC -3’; Template strand: 5’ – GTGCTGCCGAAGCGAGCACGACTCAAGGGTTACTGTCACACCTATTTTAAGCCCTTCAATCAAATCATCTTTTGGTTAGGATAACTTATGGTCGGTTT -3’) with 1pmol of labelled DNA incubated with 2-14pmol of SNAPc^FL^ or SNAPc^mini^ in 20mM HEPES pH 8.0, 250mM NaCl, 5% (v/v) glycerol at 4°C for 80mins with a total reaction volume of 20µl. In each case, the binding reactions were resolved using a 4% polyacrylamide gel (37.5:1 acrylamide/bis-acrylamide, 10% (v/v) glycerol, Tris-borate EDTA (TBE) 1X) run for 1h at 90V in 0.5X TBE running buffer. Gels were directly visualized via the Cy5 label using the BioRad ChemiDoc imaging system.

### Mass Photometry

Mass photometry experiments were performed essentially as described in Sonn-Segev et al., 2020.^77^ Standard microscope coverslips were cleaned with a Diener Pico Plasma Cleaner, silicon buffer gaskets were attached to the glass slides, and the slides were then mounted on a Refeyn OneMP mass photometer (Refeyn). For measurement, protein name samples were diluted to 10 nM in composition of the buffer immediately before measurement. The gasket was filled with 15 ul buffer and the focal plane was automatically estimated. Proteins were diluted 1:10 into the buffer-filled gasket, to a final concentration of 1 nM. A 60 second movie was then recorded through AcquireMP (Refeyn). Data was processed using the DiscoverMP program. Contrast-to-mass calibration was performed with several known-mass proteins. Mass distributions were plotted with DiscoverMP and mean mass peaks determined by Gaussian fitting.

## Data Availability

The Cryo-EM density reconstructions and final models were deposited in the Electron Microscopy Data Base (EMDB) under accession number 50730 (OC1^FL^), 50731 (OC2^FL^), 50732 (MC^FL^), 50733 (OC1^mini^) and 50734 (OC2^mini^). The corresponding model coordinates were deposited in the Protein Data Bank (PDB) under accession codes 9FSO (OC1^FL^), 9FSP (OC2^FL^), 9FSQ (MC^FL^), 9FSR (OC1^mini^) and 9FSS (OC2^mini^). The SNAPc-DNA crosslinking mass spectrometry proteomics data have been deposited to the ProteomeXchange Consortium via the PRIDE^78^ partner repository with the dataset identifier PXD053341 and 10.6019/PXD053341.

## Acknowledgements

We thank all members of the Vannini group for discussion throughout this study. We thank Alessandro Scardua and Janine Weber of the Biomass Production Unit of the National Facility for Structural Biology at Human Technopole for large-scale biomass production, technical support and assistance. Additionally, we thank Sebastiano Pasqualato and Nataliya Danilenko of the Biophysics Unit of the National Facility for Structural Biology at Human Technopole for technical support with biophysical measurements. We thank Paolo Swuec and Simona Sorrentino of the Cryo-Electron Microscopy Unit of the National Facility for Structural Biology at Human Technopole for technical support and assistance with Cryo-EM sample preparation and data collection. We also thank Daniele Colombo for technical assistance with data processing software.

## Author Contributions

A.D.M. and C.W.M. designed, supervised and performed genome-editing experiments, creating the stable POLR1C-Strep-mCherry-P2A HEK293T cell line. T.K. performed cloning for protein expression. S.Z.S., E.P.R. and V.C. performed large-scale purification of human Pol III and transcription factors, carried out biophysical characterisation, performed PIC assembly and purification and prepared samples for EM and crosslinking MS analysis. S.Z.S., E.P.R. and T.N.P. carried out Cryo-EM sample preparation, screening, sample collection and preliminary evaluation of the data. S.Z.S. and E.P.R. performed cryo-EM data processing and analysed the data. A.G. prepared samples for crosslinking MS analysis, collected data and analysed the results. A.G and E.P.R. interpreted the crosslinking MS data. S.Z.S., E.P.R. and A.V. wrote the manuscript with input from all authors. A.V. and C.W.M. supervised the research.

## Notes

### Competing Interest Statement

The authors have declared no competing interest.

