## Supplementary Data for "Structural insights into distinct mechanisms of RNA polymerase II and III recruitment to snRNA promoters"

### Extended Data Figures

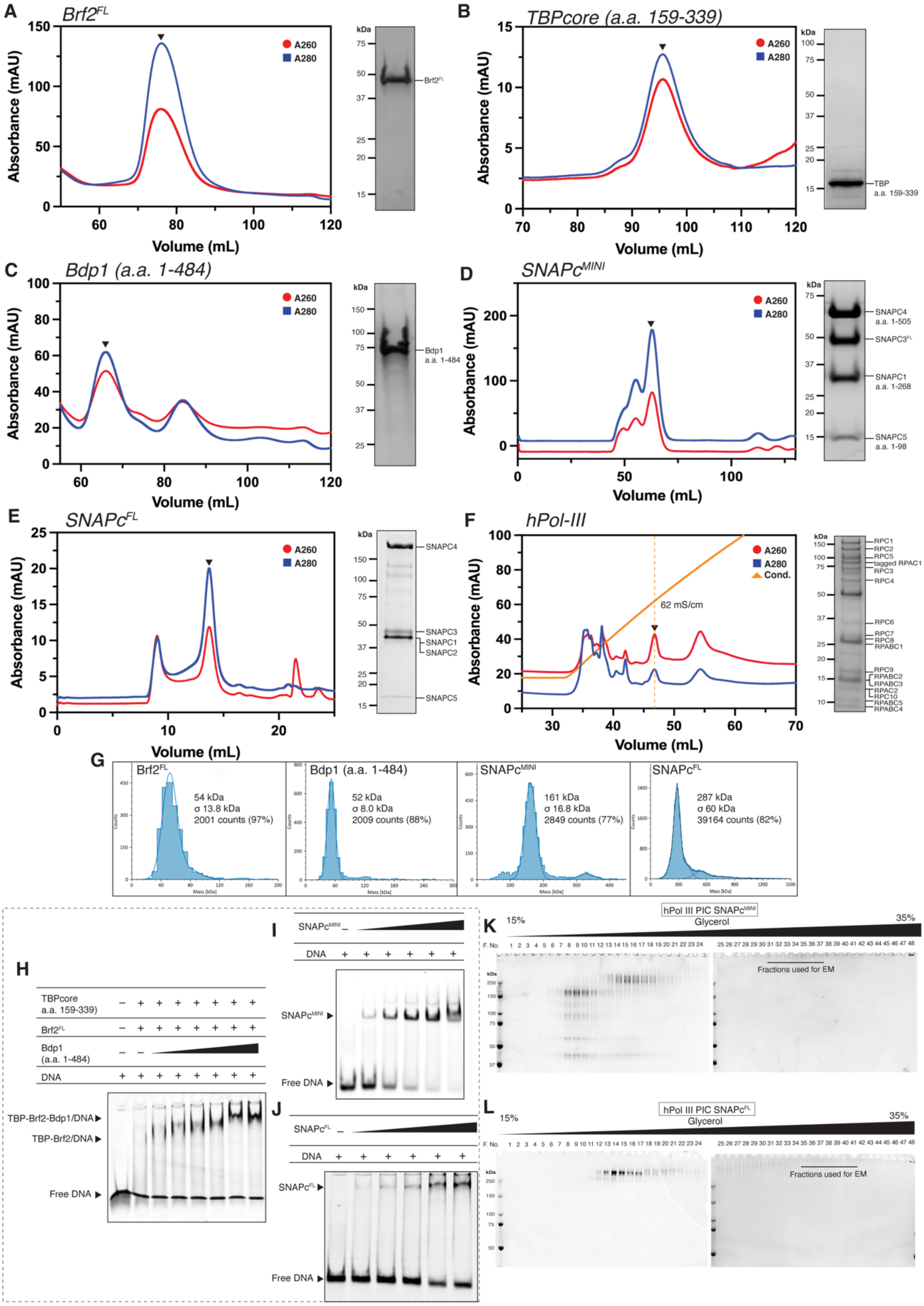


**Extended Data Figure 1 –** Protein purification and PIC complex assembly. Shown are size exclusion chromatography traces (*left*) and SDS-PAGE analysis of a representative peak fraction (*right*) for purified (a) Brf2, (b) $TBP_{core}$, (c) $Bdp1^{1-484}$, (d) $SNAPc^{mini}$, (e) $SNAPc^{FL}$ and an ResourceQ profile of (f) purified endogenous human Pol III with SDS-PAGE of the peak fraction (marked). (g) Mass photometry molecular mass determination was used to assess protein purity and oligomeric state for Brf2, $Bdp1^{1-484}$, $SNAPc^{mini}$ and $SNAPc^{FL}$. DNA binding competency of the purified transcription factor was assessed via electromobility shift assay (EMSA) for (h) $TBP_{core}$, (i) $SNAPc^{mini}$ and (j) $SNAPc^{FL}$. Following assembly of all PIC components, PICs were purified and simultaneously crosslinked using GraFix gradients. Each fraction was assessed using SDS-PAGE following PIC formation with (k) $SNAPc^{mini}$ and (l) $SNAPc^{FL}$. Fractions subject to cryo-EM analysis are highlighted.

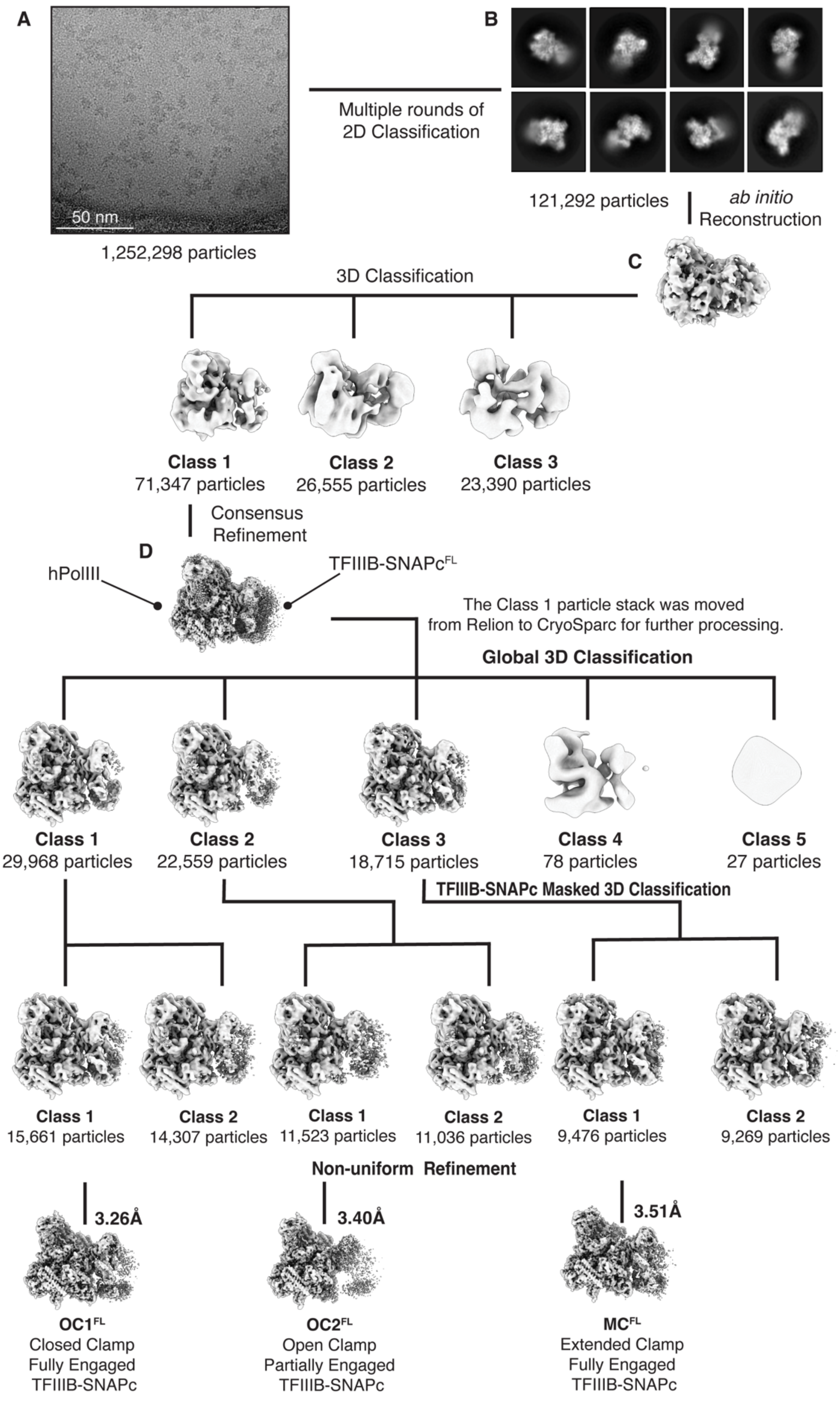


**Extended Data Figure 2 –** SNAPc$^{FL}$-PIC cryo-EM data processing. (a) Representative micrograph (from 17 291 micrographs). (b) Representative 2D class averages from 121 292 particles used subsequently for *ab initio* model building. (c) Particles were subject to 3D classification into 3 classes in cryosparc, with a single class (71 347 particles) corresponding to the PIC. (d) This was subject to global consensus refinement, with the resulting map and particles exported to RELION and subjected to a second round of 3D classification into 5 classes. (e) Classes 1, 2 and 3 represented distinct complex structures and were subject to a third round of 3D classification into 2 classes per state focused on the TFIIIB:SNAPc$^{FL}$ module. (f) A single class was refined for each state, yielding a final refined map for OC1$^{FL}$, OC2$^{FL}$ and MC$^{FL}$ at 3.26Å, 3.40Å and 3.51Å resolution, respectively.

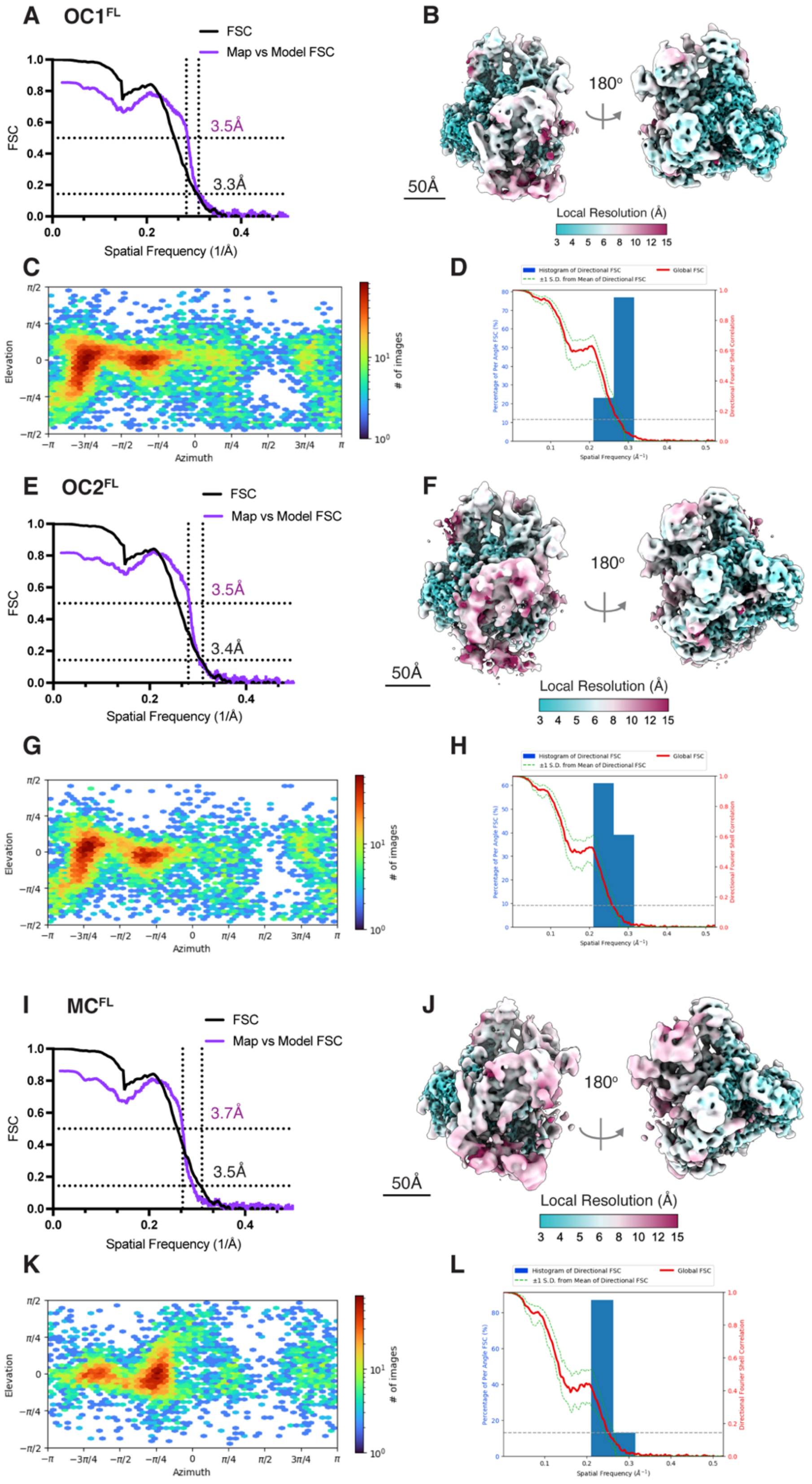


**Extended Data Figure 3 –** Resolution and density isotropy assessment of SNAPc$^{FL}$-PIC cryo-EM maps. (a, e, i) Fourier Shell correlation (FSC) of the OC1$^{FL}$, OC2$^{FL}$, MC$^{FL}$ maps (black) with the map to model FSC (purple) overlain. Resolutions are reported at the 0.143 and 0.5 criterion for the map and map to model FSC respectively. (b, f, j)

Local resolution estimation of the $OC1^{FL}$, $OC2^{FL}$, $MC^{FL}$ maps, each region of the map is filtered according to the reported local resolution. (c, g, k) Orientation distribution of particles in the $OC1^{FL}$, $OC2^{FL}$, $MC^{FL}$ reconstructions. (d, h, l) 3D FSC assessment of map isotropy, showing the global FSC (red) and 1 standard deviation around the mean (SD, green) with the 0.143 criterion marked with a dashed line. The superimposed histogram reports the percentage of voxels identified at each spatial frequency in the FSC (blue).

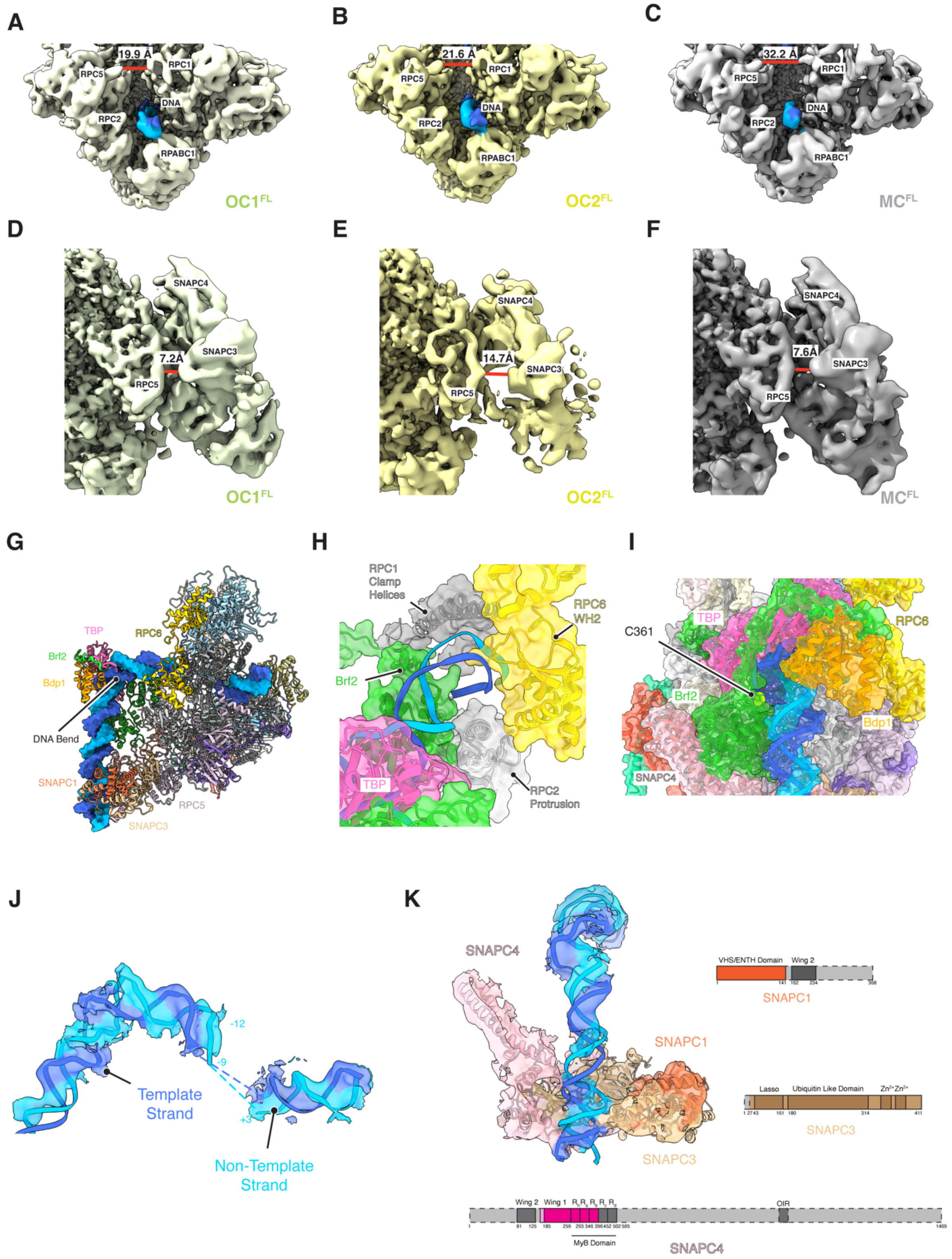


**Extended Data Figure 4 –** TFIIIB:SNAPc and DNA Binding in the SNAPc$^{FL}$-PIC **structure.** RPC1 clamp and TFIIIB: SNAPc movements observed in the SNAPc$^{FL}$-PIC cryo-EM maps. Shown are the RPC1 clamp widths measured for the (a) OC1$^{FL}$, (b) OC2$^{FL}$ and (c) MC$^{FL}$ reconstructions, with subunits labelled. Also measured were the observed TFIIIB:SNAPc movements relative to the bound polymerase, with SNAPC3-RPC5 distances measured for (d) OC1$^{FL}$, (e) OC2$^{FL}$ and (f) MC$^{FL}$. (g) The path of the bound DNA is shown in surface in the context of the PIC (ribbon) with the TBP-induced DNA bend highlighted. (h) The upstream DNA entry site into Pol III. Labelled are the subunits forming a stabilising 'channel' which feeds the DNA into the polymerase active site. (i) Highlighted in yellow is the C361 residue of Brf2, previously identified as important in the redox-mediated disassembly of the TFIIIB complex. (j) The opened DNA observed within the polymerase active site. Template and non-template strands are labelled. The non-template strand is labelled at -12 (at the site of DNA opening) and -9 (the end of the built upstream strand). (k) General features of the bound SNAPc module. Shown is the SNAPc structure fitted to cryo-EM density coloured by subunit, with corresponding diagrams highlighted (via coloured regions) the sequence portion of each subunit observed in the cryo-EM structure. Displayed in all cases is the $OC1^{FL}$ cryo-EM density.

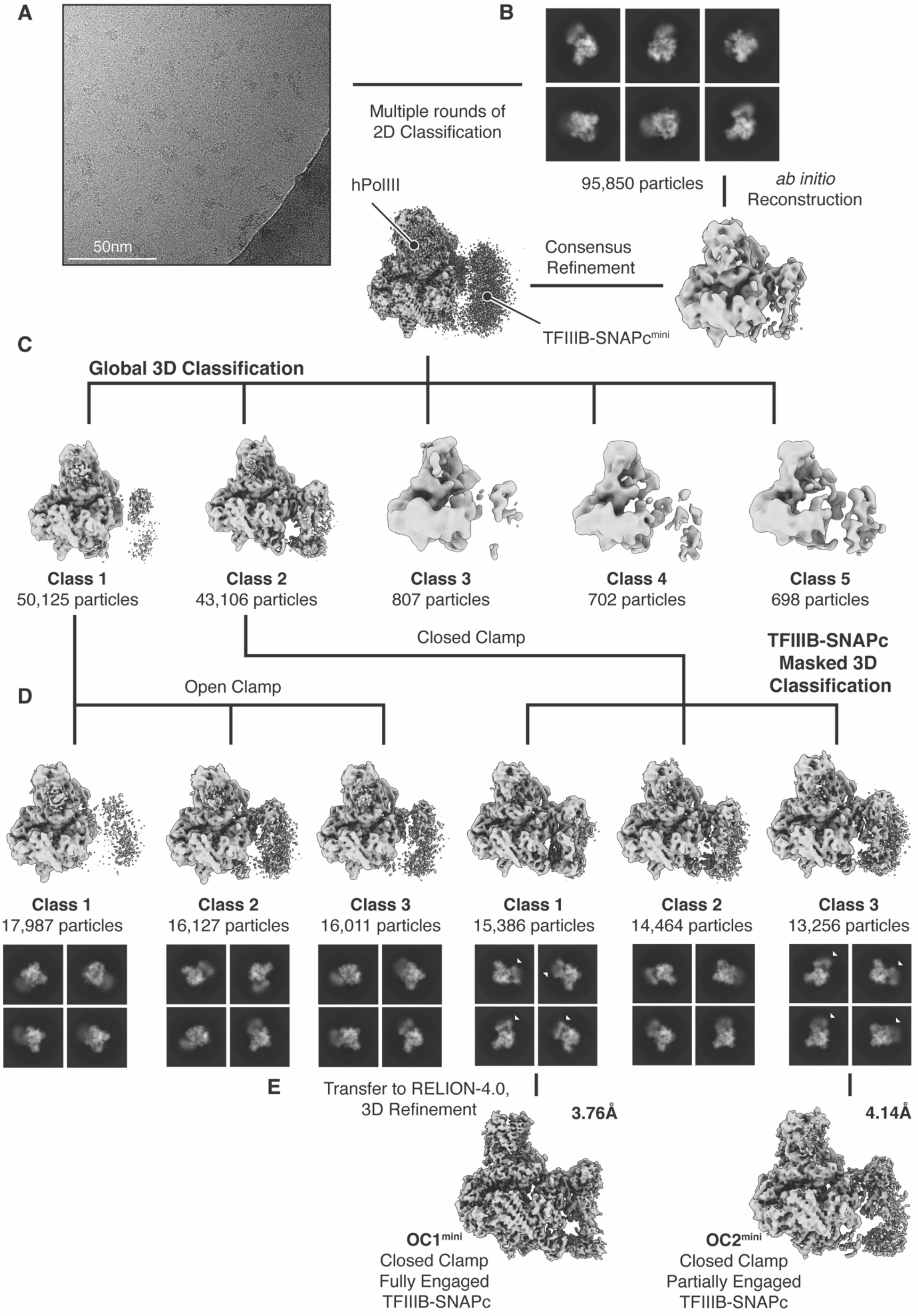


**Extended Data Figure 5 –** SNAPc$^{mini}$-PIC cryo-EM data processing. (a) Representative micrograph (from 19 060 micrographs). (b) Representative 2D class averages from 95 850 particles used subsequently for *ab initio* model building and consensus refinement. (c) Particles were subject to 3D classification into 5 classes in cryosparc, with 2 classes identified which represented the open RPC1 clamp (50 125 particles) and closed RPC1 clamp (43 106 particles) PIC. (d) Each identified PIC structure was subject to focused 3D classification around the TFIIIB:SNAPc density into 3 classes, which were also analysed by 2D classification. Only the closed clamp classes produced high quality density for the TFIIIB:SNAPc module, shown by the white arrows in subsequent 2D classes. Classes 1 and 3 represented closed clamp PICs with fully engaged (class 1) and partially engaged (class 3) TFIIIB:SNAPc. (e) These were transferred to RELION for 3D refinement, local resolution estimation and filtering to produce 2 maps named $OC1^{mini}$ and $OC2^{mini}$ at 3.8 and 4.1Å resolution, respectively.

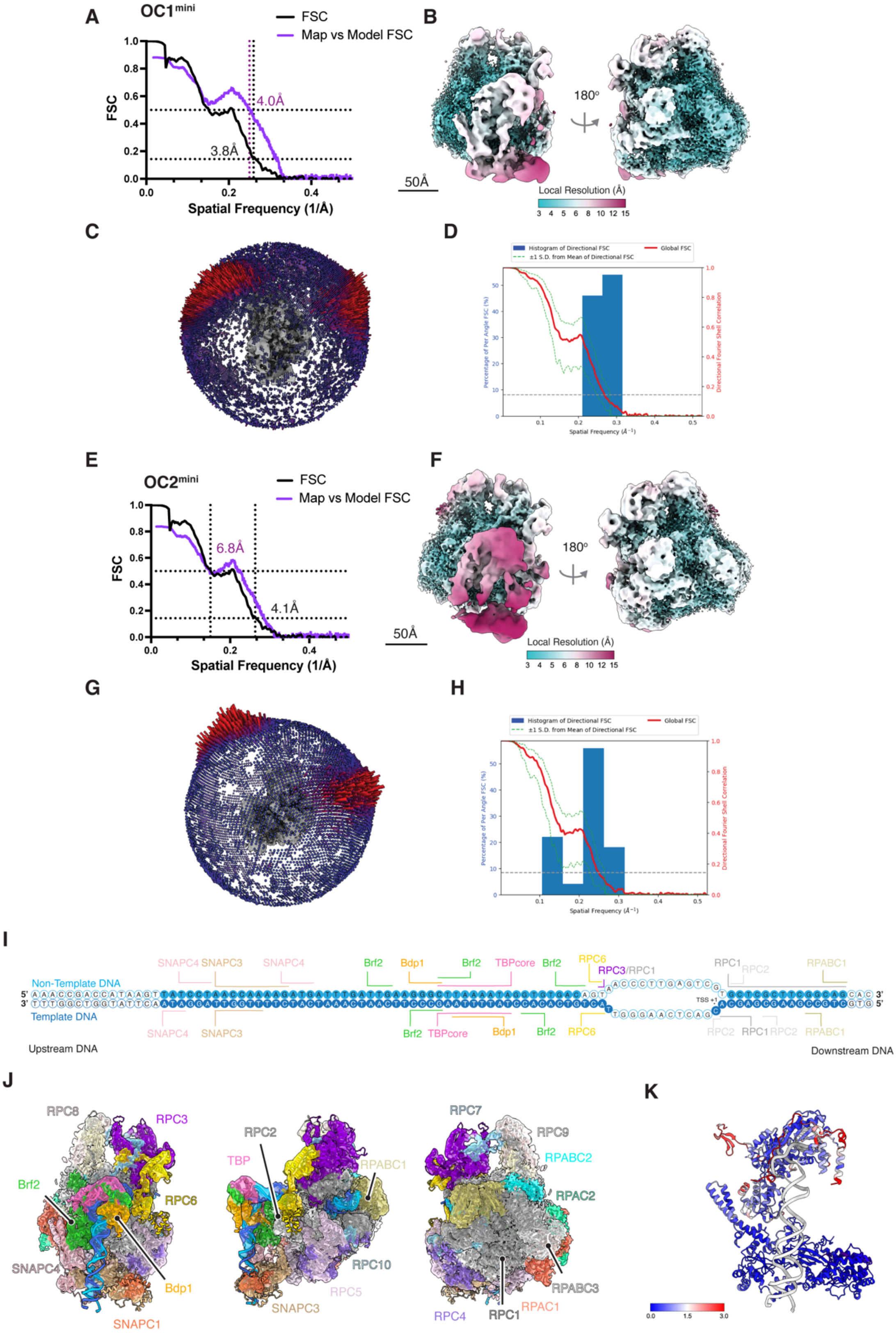


**Extended Data Figure 6 -** Resolution and density isotropy assessment of SNAPc$^{mini}$-PIC cryo-EM maps with comparison to SNAPc$^{FL}$-PIC. Resolution and density isotropy assessment of SNAPc$^{mini}$-PIC cryo-EM maps. (a, e) Fourier Shell correlation (FSC) of the OC1$^{mini}$ and OC2$^{mini}$, maps (black) with the map to model FSC (purple) overlain. Resolutions are reported at the 0.143 and 0.5 criterion for the map and map to model

FSC respectively. (b, f) Local resolution estimation of the OC1$^{mini}$ and OC2$^{mini}$ maps, each region of the map is filtered according to the reported local resolution. (c, g) Orientation distribution of particles in the OC1$^{mini}$ and OC2$^{mini}$ reconstructions. (d, h) 3D FSC assessment of map isotropy, showing the global FSC (red) and 1 standard deviation around the mean (SD, green) with the 0.143 criterion marked with a dashed line. The superimposed histogram reports the percentage of voxels identified at each spatial frequency in the FSC (blue). (i) DNA nucleotides modelled in the OC1$^{mini}$ structure are denoted a solid circles, with template and non-template strands indicated. All nucleotides are numbered according to the TSS in the human U6-2 gene. Protein-DNA interactions are labelled with the interacting protein identified. Observed DNA regions and protein contact sites were identical in both OC1$^{mini}$ and OC2$^{mini}$ reconstructions. (j) OC$^{mini}$ locally filtered cryo-EM density map with fitted structural model shown in ribbon representation. The cryo-EM density is coloured according to the fitted subunit. (k) The SNAPc$^{mini}$ structure resolved in the SNAPc$^{mini}$ PIC. Residues are shown coloured according to RMSD with the SNAPc structure observed within the SNAPc$^{FL}$-PIC.

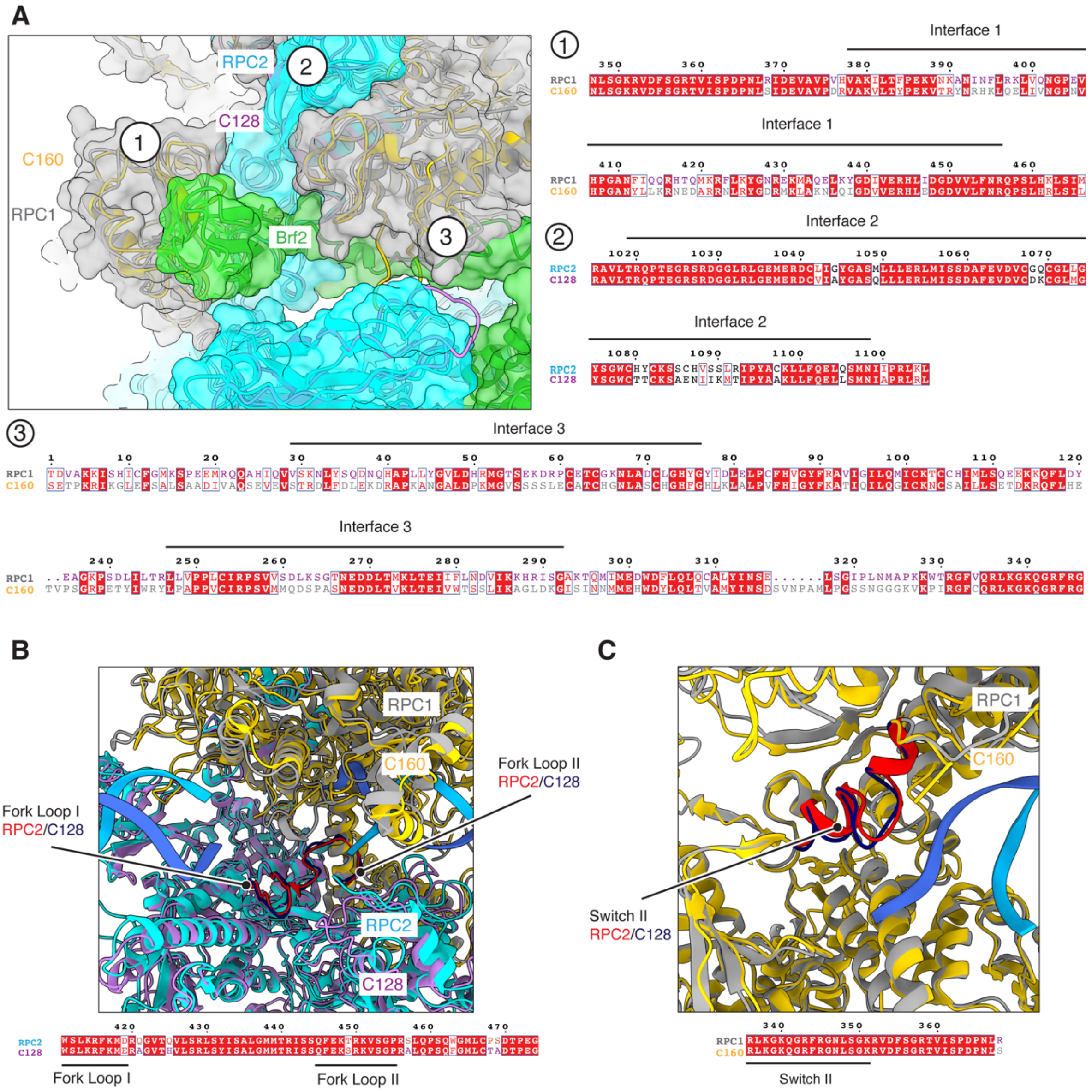


**Extended Data Figure 7 –** Structural comparison of human and yeast snRNA PICs. (a) Comparison of the human and yeast dock domain. Shown is a structural alignment between human and yeast dock domain subunits (*left*). Each observed dock domain-Zn ribbon interface is labelled (1-3) with corresponding sequence alignment for each interface displayed (*right and below*). Conserved residue identities are highlighted in red with sequence similarities outlined in blue. (b) Structural alignment of human and yeast catalytic site regions centred on fork loops I and II (labelled in red and blue for human and yeast structures respectively). Shown below is the sequence alignment for these regions. (c) Structural (*above*) and sequence (*below*) alignment of the human and yeast switch II motif (shown in red and blue respectively). In all cases, the human OC$^{FL}$ is compared to the yeast OC-PIC structure (PDB: 6eu0).

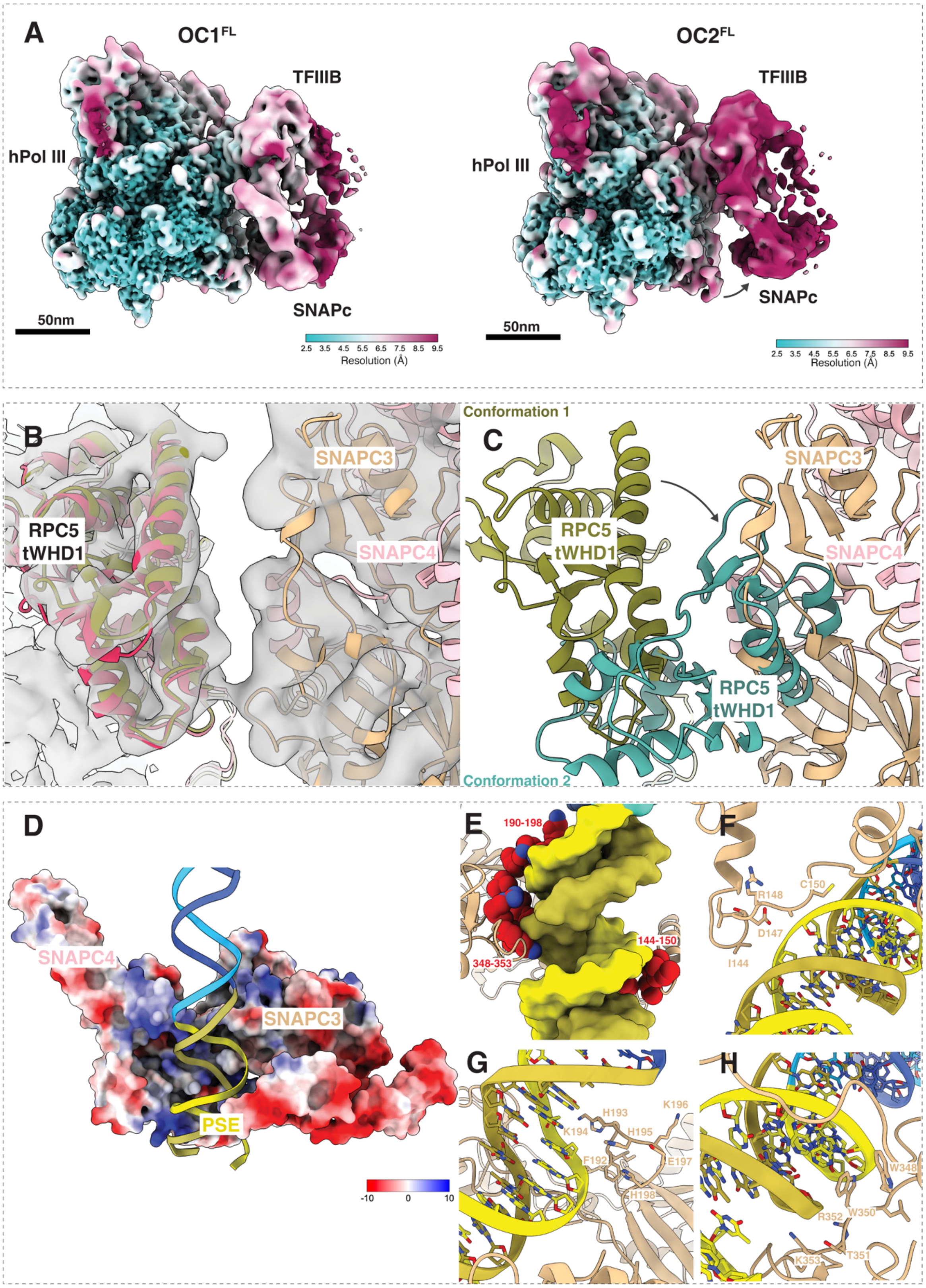


**Extended Data Figure 8** – SNAPc binding interfaces observed in PIC. (a) Local resolution estimation for both $OC1^{FL}$ and $OC2^{FL}$ highlighting the loss of local resolution of the TFIIIB:SNAPc module upon disassociation from the RPC5 C-terminal region. (b) Alignment of RPC5 tWHD1 from the PIC (green) and elongating complex (red, PDB:7ae1) showing the equivalent conformation despite the binding of SNAPC3 in the

PIC. (c) Superposition of the 2 conformations observed in the Pol III elongating complex (PDB:7ae1) onto the PIC. Whilst conformation 1 (green) is competent for assembly into the PIC, the second observed conformation (teal) induces several steric clashes with the nearby SNAPC3 subunit. (d) SNAPc binding to the PSE. Shown is the built SNAPc module in the PIC rendered according to colombic surface charge, the PSE is highlighted in yellow. (e) Main interacting regions of SNAPC3 (red) binding to the minor groove of the PSE (yellow, surface representation). Representative SNAPC3-PSE interactions from each interaction region are shown in each panels (f-h).

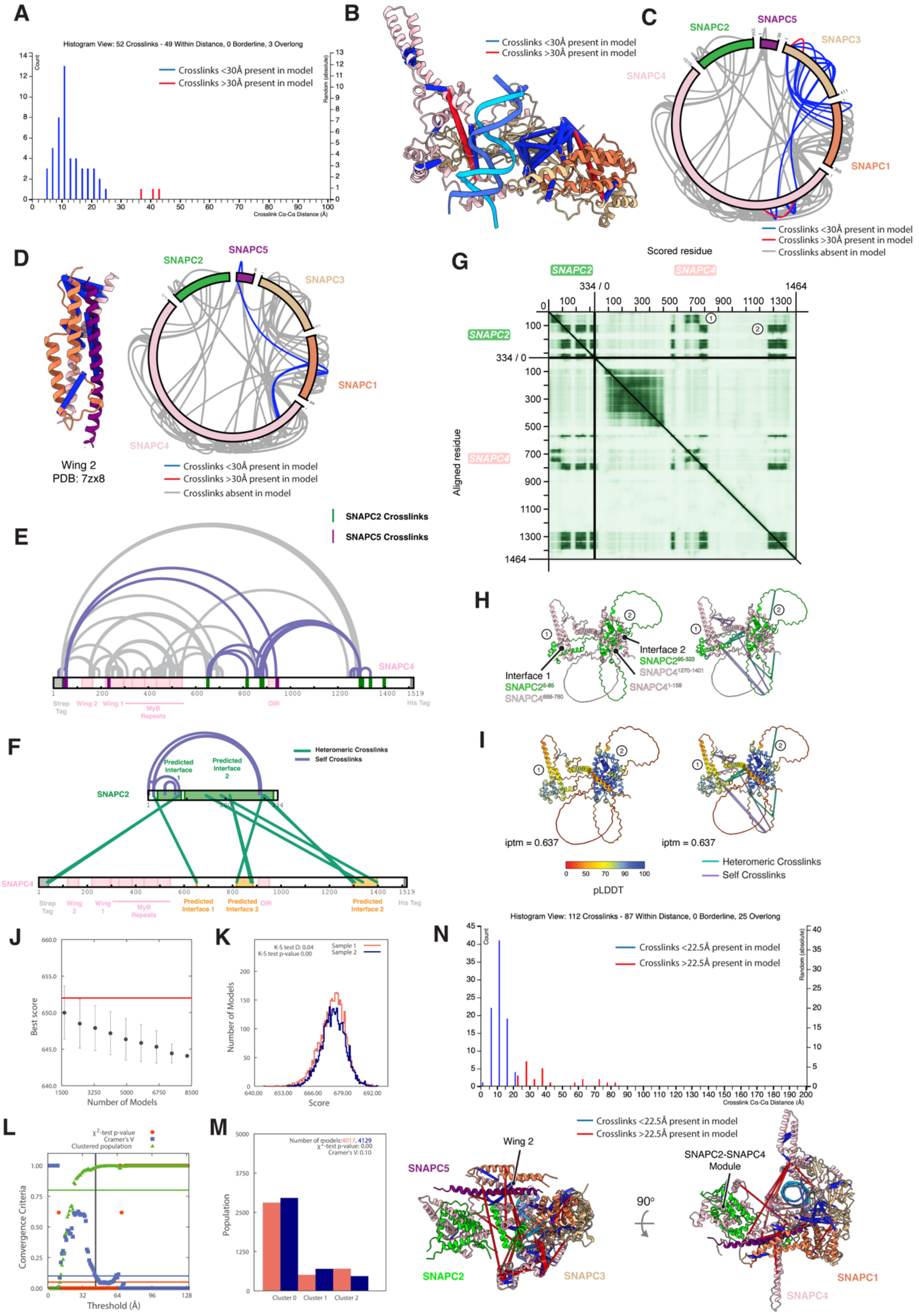

**Extended Data Figure 9** – Sulfo-SDA crosslinking mass spectrometry and alphafold provide insight into SNAPC2 and SNAPC5 localisation. (a) Histogram of all crosslinks observed within the OC$^{FL}$ SNAPc structure, with crosslinks coloured according to satisfaction (blue) and violation (red) of the 30Å distance restraint. (b) Mapping of the observed crosslinks in SNAPc according to distance restraint violation, also visualized in the context of all crosslinks identified in the complex (c). (d) Mapping of SNAPc-DNA crosslinks onto the wing 2 structure from the Pol II U1-snRNA PIC (PDB:7zx8). Crosslinks are coloured according to distance restraint violation and shown mapped directly onto the wing 2 structure (*left*) and in the context of all identified SNAPc-DNA crosslinks (*right*). (e) Self crosslinks observed within the SNAPC4 subunit. Highlighted are SNAPC2 (green) and SNAPC5 (purple) crosslinks, with SNAPC4 self-crosslinks observed linking these contact sites shown in lilac. (f) Mapping the alphafold SNAPC2-SNAPC4 interface predictions onto the crosslinking data revealed agreement. Alphafold predicted interfaces in SNAPC2 (green) and SNAPC4 (orange) are shown with heteromeric crosslinks between the proteins shown in green. (g) Shown is the Predicted Aligned Error (PAE) plot for the alphapulldown prediction of the SNAPC2-SNAPC4 interaction site. The two identified interfaces are labelled. (h) The alphapulldown SNAPC2-SNAPC4 complex prediction, showing both identified interfaces (*left*) with the identified crosslinks mapped onto the structure (*right)*. (i) The SNAPC2-SNAPC4 complex prediction rendered according to predicted local distance difference test (pLDDT) with crosslinks mapped (*right*). Reported also is the interface predicted template modelling score (iptm) of 0.637 for the prediction. IMP modelling of mobile regions in the SNAPc-DNA structure. (j) Shown is the convergence of the structure score for random selections of the 8259 clustered models included in the analysis. All structural scores are similar and within error irrespective of the number of models computed. (k) Distribution of scores for model sample populations 1 and 2 encompassing the 8259 models, with 4017 and 4129 models in populations 1 and 2 respectively. The Kolmogorov-Smirnov (K-S) test shows that the difference between the two distributions of structure scores is small (with a D=0.04) and insignificant (with a p-value of 0) and so both score distributions for each population are highly similar. (l) Determination of global sampling precision using RMSD of computed models for all defined rigid bodies. Statistical significance is calculated via $c^2$ p-values (red points), Cramer's V value (blue squares) and the population of structures in sufficiently large clusters (green triangles) The vertical dotted grey line indicates the RMSD which defines the global sampling precision of 44 Å. (m) Sample populations in each computed cluster obtained by clustering via an RMSD threshold of 44Å. Cluster 0 represents the dominant cluster containing 70% of all calculated structures (cluster 0 sample 1 = 2807 models, sample 2 = 2959 models, total = 5766 models). The precision of cluster 0 is 27 Å. (n) Histogram of all crosslinks observed within the centroid model for cluster 0, with crosslinks coloured according to satisfaction (blue) and violation (red) of the 22.5Å distance restraint used during IMP simulations (*above*) with mapping of the sulfo-SDA crosslinks onto the centroid cluster model (cluster 0) coloured according to satisfaction (blue) and violation (red) of the 22.5 Å distance restraint (*below*).

**A**

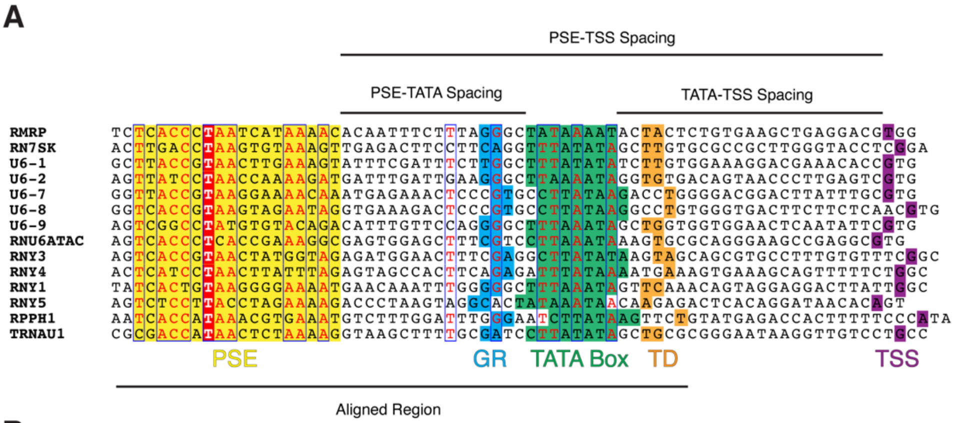


**B**

| | PSE-TATA Box Distance (nt) | TATA Box-TSS Distance (nt) | PSE-TSS Distance (nt) |
|---|---|---|---|
| RMRP | 16 | 23 | 47 |
| RN7SK | 16 | 24 | 48 |
| U6_1 | 16 | 23 | 47 |
| U6_2 | 16 | 23 | 47 |
| U6_7 | 17 | 22 | 47 |
| U6_8 | 17 | 24 | 49 |
| U6_9 | 16 | 23 | 47 |
| RNU6ATAC | 16 | 22 | 46 |
| RNY3 | 17 | 24 | 49 |
| RNY4 | 17 | 23 | 48 |
| RNY1 | 16 | 24 | 48 |
| RNY5 | 15 | 23 | 46 |
| RPPH1 | 18 | 24 | 50 |
| TRNAU1 | 16 | 24 | 48 |

**Extended Data Figure 10** – Alignment of Pol III class III promoters. (a) Multiple sequence alignment of all pol III class III promoters. The PSE and TATA boxes are highlighted with yellow and green boxes, respectively. The conserved GR and TD motifs found flanking the TATA box are highlighted in blue and orange respectively, with the transcriptional start site (TSS) highlighted in purple. Conserved nucleotides are denoted in red. (b) Table showing the calculated TATA-PSE, TATA-TSS and PSE-TSS spacings for each Pol III class III promoter

### Extended Data Tables

| OC1$^{FL}$ - OC2$^{FL}$ | | |
|---|---|---|
| **Subunit** | **RMSD (Å)** | **No. Residue Pairs Analysed** |
| **Brf2** | 0.433 | 361 |
| **Bdp1** | 0.571 | 78 |
| **TBPcore** | 0.419 | 178 |
| **SNAPC1** | 0.152 | 141 |
| **SNAPC3** | 0.155 | 385 |
| **SNAPC4** | 0.272 | 242 |
| **OC1$^{FL}$ - MC$^{FL}$** | | |
| **Subunit** | **RMSD (Å)** | **No. Residue Pairs Analysed** |
| **Brf2** | 0.487 | 361 |
| **Bdp1** | 0.599 | 78 |
| **TBPcore** | 0.421 | 178 |
| **SNAPC1** | 0.192 | 141 |
| **SNAPC3** | 0.227 | 385 |
| **SNAPC4** | 0.303 | 242 |
| **OC1$^{FL}$ - OC1$^{mini}$** | | |
| **Subunit** | **RMSD (Å)** | **No. Residue Pairs Analysed** |
| **Brf2** | 0.95 | 351 |
| **Bdp1** | 1 | 78 |
| **TBPcore** | 0.77 | 178 |
| **SNAPC1** | 0.39 | 141 |
| **SNAPC3** | 0.37 | 385 |
| **SNAPC4** | 0.53 | 242 |
| **OC1$^{FL}$ - OC1$^{mini}$ (PDB: 7xur)** | | |
| **Subunit** | **RMSD (Å)** | **No. Residue Pairs Analysed** |
| **SNAPC1** | 0.95 | 133 |
| **SNAPC3** | 1.17 | 247 |
| **SNAPC4** | 1.28 | 101 |
| **H.s. Brf2 C-Cyclin Comparisons** | | |
| **Subunit** | **RMSD (Å)** | **No. Residue Pairs Analysed** |
| **H.s Brf2 v H.s. Brf1** | 1.287 | 108 |
| **H.s. Brf2 v S.c. Brf1** | 1.2 | 102 |
| **H.s. Brf2 v TFIIB** | 1.425 | 94 |

**Extended Data Table 1 –** Pairwise RMSD comparison between Pol III PIC components. Models compared are highlighted in bold with the subunit, RMSD and number of residues included each pairwise comparison reported.

| Protein | Residue Range | Residues Per Bead | Rigid Body Number | Structural Model Source (PDB Identifier/this study) |
|---|---|---|---|---|
| **SNAPC1** | 1-141 | 10 | 1 | This Study, OCFL structure |
| **SNAPC1** | 142-161 | 10 | 4 | - |
| **SNAPC1** | 162-234 | 10 | 4 | PDB: 7zx8 |
| **SNAPC1** | 235-368 | 20 | 4 | - |
| **SNAPC2** | 1-28 | 10 | 2 | - |
| **SNAPC2** | 29-87 | 10 | 2 | This Study, SNAPC2-SNAPC4 Alphafold Multimer |
| **SNAPC2** | 88-96 | 10 | 3 | - |
| **SNAPC2** | 97-160 | 10 | 3 | This Study, SNAPC2-SNAPC4 Alphafold Multimer |
| **SNAPC2** | 161-200 | 10 | 3 | - |
| **SNAPC2** | 201-271 | 10 | 3 | This Study, SNAPC2-SNAPC4 Alphafold Multimer |
| **SNAPC2** | 272-303 | 10 | 3 | - |
| **SNAPC2** | 304-334 | 10 | 3 | This Study, SNAPC2-SNAPC4 Alphafold Multimer |
| **SNAPC3** | 1-26 | 10 | 1 | - |
| **SNAPC3** | 27-411 | 10 | 1 | This Study, OCFL structure |
| **SNAPC4** | 1-81 | 20 | 4 | - |
| **SNAPC4** | 83-124 | 10 | 4 | PDB: 7zx8 |
| **SNAPC4** | 125-141 | 20 | 4 | - |
| **SNAPC4** | 142-377 | 10 | 1 | This Study, OCFL structure |
| **SNAPC4** | 378-383 | 20 | 1 | - |
| **SNAPC4** | 384-502 | 10 | 1 | PDB: 7xur |
| **SNAPC4** | 503-665 | 20 | 1 | - |
| **SNAPC4** | 666-759 | 10 | 2 | This Study, SNAPC2-SNAPC4 Alphafold Multimer |
| **SNAPC4** | 760-765 | 20 | 2 | - |
| **SNAPC4** | 766-819 | 10 | 3 | This Study, SNAPC2-SNAPC4 Alphafold Multimer |
| **SNAPC4** | 820-1263 | 20 | 3 | - |
| **SNAPC4** | 1264-1394 | 10 | 3 | This Study, SNAPC2-SNAPC4 Alphafold Multimer |
| **SNAPC4** | 1395-1469 | 20 | 3 | - |
| **SNAPC5** | 1-52 | 10 | 4 | PDB: 7zx8 |
| **SNAPC5** | 53-98 | 10 | 4 | - |

| | | | | |
|---|---|---|---|---|
| **Template Strand, DNA** | 1-20 | 10 | 1 | This Study, OCFL structure |
| **Non-template Strand, DNA** | 1-20 | 10 | 1 | This Study, OCFL structure |

**Extended Data Table 2 –** Coarse graining representations for the full length SNAPc complex in IMP. Shown are the modelled proteins, divided by regions corresponding to defined structures or flexible beads. Each region belongs to a defined rigid body (numbered 1-5) with rigid body 1 representing the fixed region derived from the $OC^{FL}$ model.
